# On the fitness of informative cues in complex environments

**DOI:** 10.1101/2020.04.28.066571

**Authors:** Fabrizio Mafessoni, Michael Lachmann, Chaitanya S. Gokhale

## Abstract

To be able to deal with uncertainty is of primary importance to most living organisms. When cues provide information about the state of the environment, organisms can use them to respond flexibly. Life forms have evolved complex adaptations and sensory mechanisms to use these environmental cues and extract valuable information about the environment. Previous work has shown a theoretical limit to the amount of fitness benefit possible to be extracted from the cues. We show that the previously used information theoretical approaches can be generalised to scenarios involving any potential relationship between the number of possible phenotypes and environmental states. Such cases are relevant when physiological constraints or complex ecological scenarios lead to the number of environmental states exceeding potential phenotypes. We illustrate cases in which these scenarios can emerge: along environmental gradients, such as geographical transects or complex environments, where organisms adopt different bet-hedging strategies, switching stochastically between phenotypes or developing intermediate ones. In conclusion, we develop an information-theoretic extensible approach for investigating and quantifying fitness in ecological studies.

## 1 Introduction

Change is a significant constant in the natural world. Organisms have to be plastic enough to ride out the variability in the environment. A potential strategy to cope with this variability is to be plastic by relying on information provided by environmental cues. An informative cue is “a feature of the world, animate or inanimate, which an animal can use to guide future actions.” [1, 2], or development. Many organisms demonstrate a plastic response to the environment, using environmental cues to modulate their phenotype [3].

However, some of the most basic adaptive mechanisms do not require informative cues. For instance, a comprehensive genetic diversity results in a better response of populations to random environmental fluctuations [4]. Another way can be for an organism to adopt a life history of “bet-hedging”. This is possible via the adoption of a generalist phenotype (conservative bet-hedging) or by randomly switching between different phenotypes, within or between generations (diversifying bet-hedging) [5, 6, 7, 8]. Hereafter we use bet-hedging as a synonym for diversifying bet-hedging.

Even without genetic variation in the population, bet-hedging enables coping with uncertain environments [9, 10, 11]: when all individuals in a population experience the same environmental state, developmental variability in the phenotypes of an individual’s offspring will ensure that at least part of its progeny will develop a proper phenotype for the current environmental state. Instances of this phenomenon can be observed in nature [12, 13, 14]. The theory of bet-hedging has a long history [15, 16, 17, 18] and hinges on the trade-off between maximizing short term fitness and reducing the adverse long term effects of its variability due to the stochasticity of the environment. Interestingly, even when cues are present but do not provide complete information, bet-hedging can still occur due to their uncertainty [19].

Previous studies quantified the fitness values of informative cues by comparing the fitness of plastic strategies relying on cues and bet-hedging strategies not relying on cues. These studies pointed to a tight relationship between evolution and information. In simple cases where the number of phenotypes equals that of environmental states, bits can quantify the evolutionary benefit of an informative cue [20]. While information theory provides a promising framework to study evolution, relatively few studies used this framework to study ecological and evolutionary phenomena [21]. In this study, we extend the information theory approach previously described by Bergstrom and Lachmann [20, 22] to examine the effects of more complex scenarios. In nature, often environmental complexity exceeds physiological and evolutionary flexibility, and the number of possible environmental states might exceed that of phenotypes. For example, the phenotypes might lie far from each other in the fitness landscape to be accessible, or physiological constraints in an organism might limit the variety of phenotypes it can develop. Furthermore, one of the available phenotypes may perform better in more than one environment. In these cases, as environmental complexity increases, the optimal strategy is more likely the one in which the same phenotypes are adopted in response to different environmental states, i.e. conservative bet-hedging occurs along with diversifying bet-hedging. These instances, in which the number of environmental states exceeds the number of phenotypes, are the focus of this study. We show that in these cases, vibrant patterns of bet-hedging emerge and that the potential fitness value of informative cues can decrease.

We first begin by reviewing the connection between Shannon and Gould information as per [20, 22], by following the calculations from Donaldson-Matasci *et al.* [22], where an organism can have as many phenotypes as the number of environmental states (represented by a square matrix). Then we generalize this analysis to any relationship between the number of phenotypes and the number of environmental states. Inequality between the number of phenotypes and environmental states (represented by a non-square matrix) is also possible. We explore representative cases of asymmetric scenarios, characterized by simple probability distributions describing the occurrence of different environmental states. In these cases, the best bet-hedging strategy depends on the probability of the environmental states in a nonlinear fashion. We discuss some examples, among which those of organisms adapted only to a range of environmental conditions, for which often increased environmental uncertainty is present at the borders of their distribution. We show that in relatively more complex scenarios, the fitness value of the informative cues is less than the mutual information between the cue and the environment. We generalize this observation to any asymmetric scenarios, showing a calculable lower bound to the fitness benefit of an informative cue under natural asymmetric conditions of complex environments.

In biology, a direct method to quantify the impact of a cue is to compare the fitness when knowing and not knowing the cue. The difference in the fitness of the strategy adopted when knowing the cue and not knowing the cue is the value of the cue measured in units of fitness gained [23, 24]. Such cues are essential across various examples: from the germination in seeds and diapause in insects to offspring clutch size control by parents. If a cue provides complete information about the predators’ presence, the prey can opt for a proper course of action. This increase in fitness measured in the information gain is **Gould information**.

A classical method to quantify the value of an environmental cue is **Shannon Information**. Consider a fair coin. The two possible outcomes “heads” (1) and “tails” (0) are equally likely with equal probabilities (*p*_0_ = *p*_1_ = 0.5). The amount of surprise which we have when we know the result of a certain coin toss event *E* is *H*(*E*) = −*p*_0_ log_2_ *p*_0_ − *p*_1_ log_2_ *p*_1_ which in this case equals 1. This is the Shannon entropy as defined in classical information theory [25, 26]. Traditional approaches in statistical physics, communication, engineering and related fields make use of the concept of ‘mutual information’. The amount by which a cue (*C*) reduces the uncertainty about the environment (*E*) is defined as mutual information (*I*(*E*; *C*)). It is measured in terms of entropy difference as

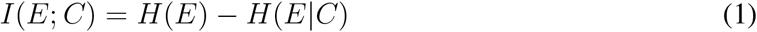

where *H*(*E*) = −Σ_*e*_ *p*_*e*_log *p*_*e*_ is the ‘entropy’ of the random variable *E* denoting the environment. Similarly *H*(*E*|*C*) = − Σ_*c*_ *p*(*e*|*c*) log *p*(*e*|*c*) is the entropy when the cue has been received. Here *p_c_* is the probability of observing the cue *c* and the probability that the environment is in state *e* when cue *c* is observed is given by *p*(*e*|*c*). If a cue is not related to the environment, then the entropy remains unchanged (*H*(*E*|*C*) = *H*(*E*)) and the mutual information between the cue and the environment is zero. However for a perfect cue this probability is *p*(*e*|*c*) = 1 and hence *H*(*E*|*C*) = 0. Thus the cue reveals the environment entirely, and the mutual information is precisely equal to the entropy of the system *I*(*E*; *C*) = *H*(*E*).

Strong links exist between the two measures of the value of informative cues [20, 22, 27, 28]. Studies show that if the environmental cue is flawlessly informative, then the best approach is to bet-hedge for an intermediate probability of an adverse event. This region of probability space in which bet-hedging occurs is a function of the Shannon entropy, and the fitness value of information is bounded above by the Shannon entropy [20]. Thus an intimate connection exists between the classical information-theoretic approach and the biologically intuitive Gould information approach.

Throughout the analyses we adopt a geometric mean approach [29, 30, 22]. Whereas the arithmetic-mean does not capture the effects of environmental variance on fitness, the multiplicative nature of geometric-mean efficiently describes the effects on the growth rate as long as there are no interactions between the phenotypes, i.e. where the average fitness is frequency independent [15, 31, 32, 33]. Indeed, in a variable environment the allele having the higher geometric mean takes over the population [34, 35, 36]. Therefore, short term fitness, usually considered in evolutionary game theory for the analysis of frequency-dependent dynamics, might not be the most helpful statistic in the case of environmental variability. Instead, a geometric mean approach can be fruitfully adopted in these cases, elucidating the role of bet-hedging strategies in minimizing fitness variability in the absence of informative cues [20, 22, 8].

## 2 Model and Results

### 2.1 Symmetric case: two environmental states with two phenotypes

Traditionally bet-hedging models have focused on the example of annual plants [15]. We use it here as well, describing a representative case using simple fictitious interaction matrices. A detailed description of possible ecological interpretations follows in the discussion.

A year could be wet *e*_1_ or dry *e*_2_. Seeds of desert annuals are better off staying dormant over dry years and do better by germinating in wet years. This relationship can be represented in the form of an interaction matrix as follows:

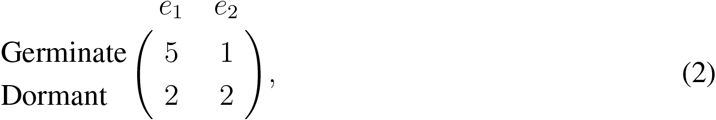

The probability that a flash flood occurs and a year is wet is *p*, while the year is dry with the probability (1 – *p*). The gain of the two phenotypes averaged over the two environments are given by:

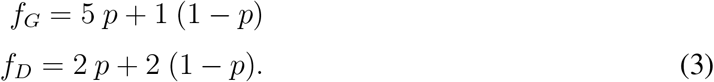

These are the average fitnesses of the two phenotypes. In the short run organisms will maximize their expected fitness by employing a strategy that maximizes its single generation expected fitness, *F* (*E*) = *max*[*f_D_, f_G_*] where *E* is a random variable representing the state of the environment (Fig. 1). The internal equilibrium of this system is at *p*^∗^ = 1/4, where the fitnesses of the two strategies are equal. Hence if the probability of a flash flood is greater than *p*^∗^ it is better to germinate else staying dormant is a safe bet:

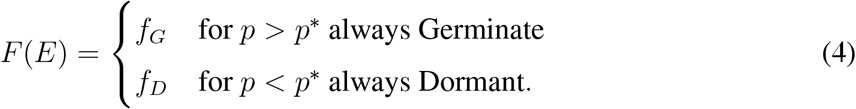

**Figure 1:**
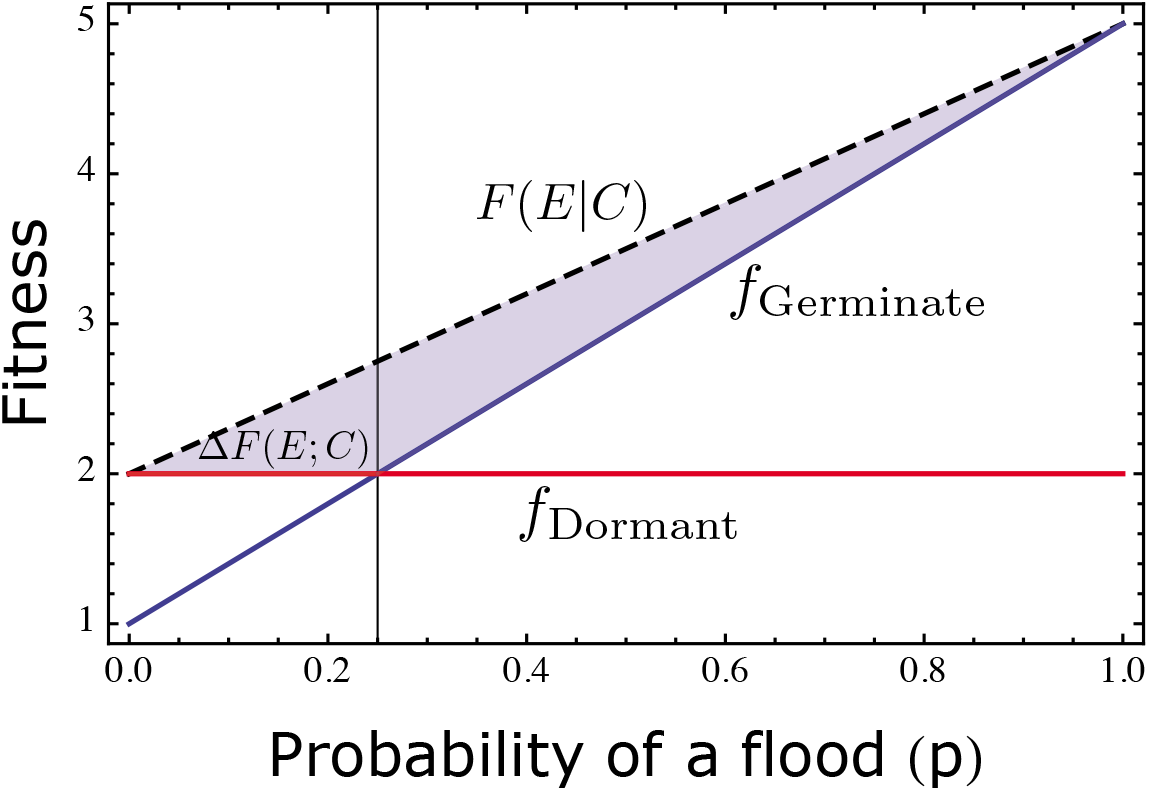
Average fitnesses of the Germinate and remain Dormant strategies as a function of the probability of a flood *p*. The internal equilibrium is given by the vertical line at *p*^∗^ = 1/4. For *p* < *p*^∗^ it pays to remain dormant while for *p* > *p*^∗^ it is better to germinate. When knowing the state of the environment exactly from the cue, a seed can obtain *F* (*E*|*C*). Hence the fitness benefit due to the cue is given by the shaded area.

In order to quantify the reduction in fitness due to environmental uncertainty, we can now suppose that the organisms might receive a cue *C*, which indicates the environmental state precisely. Therefore, then there is no confusion over choosing to germinate or not regardless of *p*. In this case the single generational expected fitness will be, *F* (*E | C*) = 5*p* + 2(1 – *p*). The value of the cue is then the difference between the fitness with the cue and without the cue:

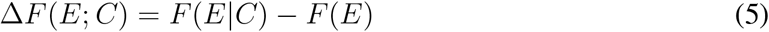

This is the result which we expect if the organism maximizes the single generational expected fitness. However, in a temporally varying environment, the most likely phenotype to fix over the long term is the one with the highest expected long-term growth rate. To approach this question, we define the frequencies with which each phenotype is adopted in a given season as *x* (Germinate) and 1 − *x* (Dormant), respectively. The mean log fitness can estimate the longterm growth rate of a given strategy, or, equivalently, the log of the geometric mean fitness [15, 37, 22, 19]. The expected long term growth rate is then given by

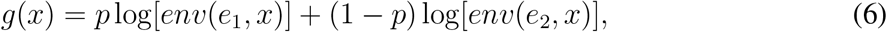

where the fitnesses in the two environments are given by:

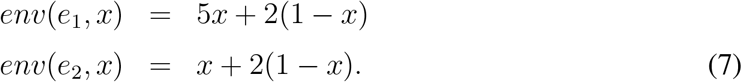

Maximizing the expected long term growth rate results in

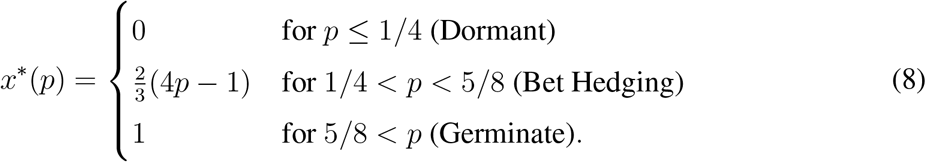

Thus, we can identify a bet-hedging region, namely a region in probability space where probabilistic switching between the phenotypes occurs. Substituting this result back in the growth rates we get

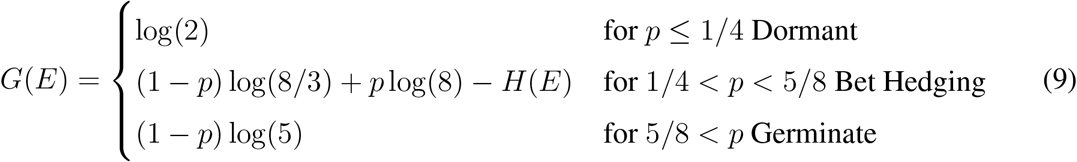

where *H*(*E*) = −*p* log(*p*)−(1−*p*) log(1−*p*) is the entropy of the random variable *E*. Making use of a perfectly informative cue the growth rate can be given by *G*(*E*|*C*) = *p* log(5) + (1 − *p*) log(2). Hence now the value of the cue (Fig. 2) is the difference between the growth rate with the cue and the one without, i.e. *G*(*E*|*C*) − *G*(*E*), and equals:

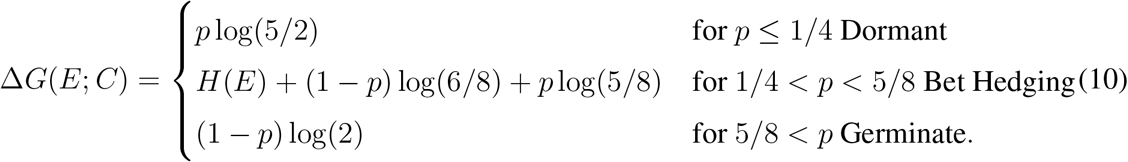

**Figure 2:**
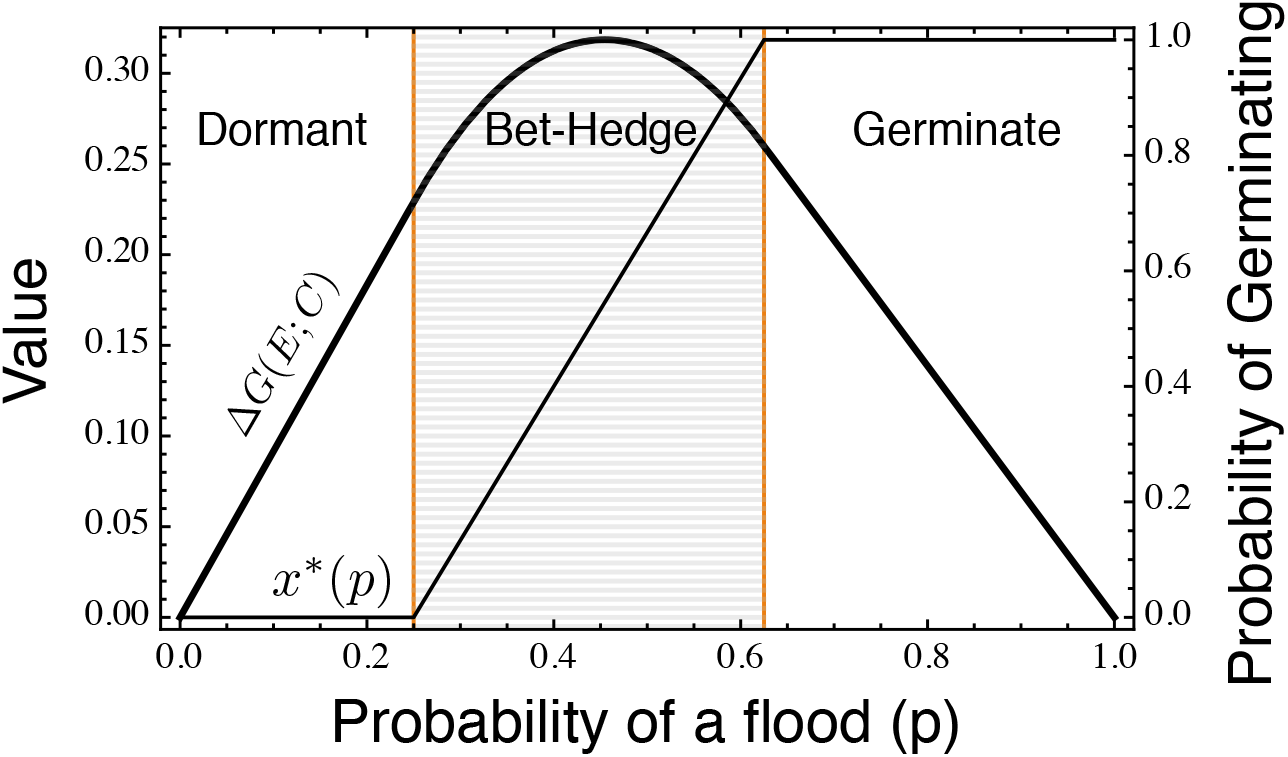
The value of the cue and the probability to germinate as a function of the probability of environmental state *e*_1_. The value of the cue ∆*G*(*E*; *C*) increases until it peaks within the area of bet-hedging. The bounds of the area of bet-hedging can be calculated analytically (see Eq. (8)). These bounds delineate the transitions in the effective probability of germinating *x*^∗^(*p*) which shifts from 0 (Dormancy) to 1 (Germination) via a linear increase within the area of bet-hedging.

Previous studies show that ∆*G*(*E*; *C*) peaks within the region of bet-hedging and is bounded by the mutual information between the cue and the environment [20]. When the cue is perfect, mutual information is simply the Shannon entropy of the environment *H*(*E*).

### 2.2 Examples with asymmetry in strategies and environments

The environment can vary in time and space. Generally, an organism is adapted only to a limited range within an environmental spectrum. Thus, including intermediate environmental states can better represent relevant environmental variability. For example, the risk of meeting a predator or the occurrence of certain climatic events can vary along both large or short ecological gradients. For instance, precipitation can vary enormously along with geographical distances. This variability further determines the distribution of plant species since for seedlings, a moderate amount of rain is preferable, while drought or extreme flooding may limit their growth.

Here we consider the different possible fitness effects of multiple environmental states, for example, seed dormancy. For simplicity, we take into account the possibility that two consecutive storms occur within one season (*e*_1_), only one (*e*_2_), or none, leading to a dry year (*e*_3_). These events occur respectively with probabilities *p*^2^, 2(1−*p*)*p* and (1−*p*)^2^. In principle, we can assume a complicated function with two variables instead of only *p*. However, as we show below, even this simple parameterisation can result in intricate bet-hedging patterns, which is sufficient to make our point.

#### 2.2.1 Adaptation to intermediates

Assuming that a single flood might provide the necessary humidity but two consecutive ones might damage the seedlings we can write down the following interaction matrix:

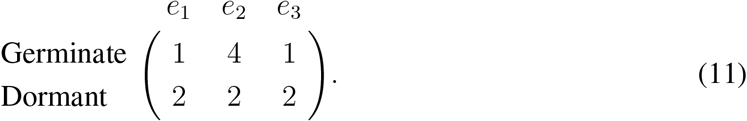

We have a non-monotonic behaviour of the value of the cue (Appendix A, Fig. 3 (a)). However, the probability of playing the ‘Always Germinate’ strategy never materialises. Instead, the seeds do best, remaining dormant close to the extreme conditions and only germinating about half the time at most when hedging their bets. The bet-hedging region in this particular case explores the mixed phenotype space between the pure phenotypes.

**Figure 3:**
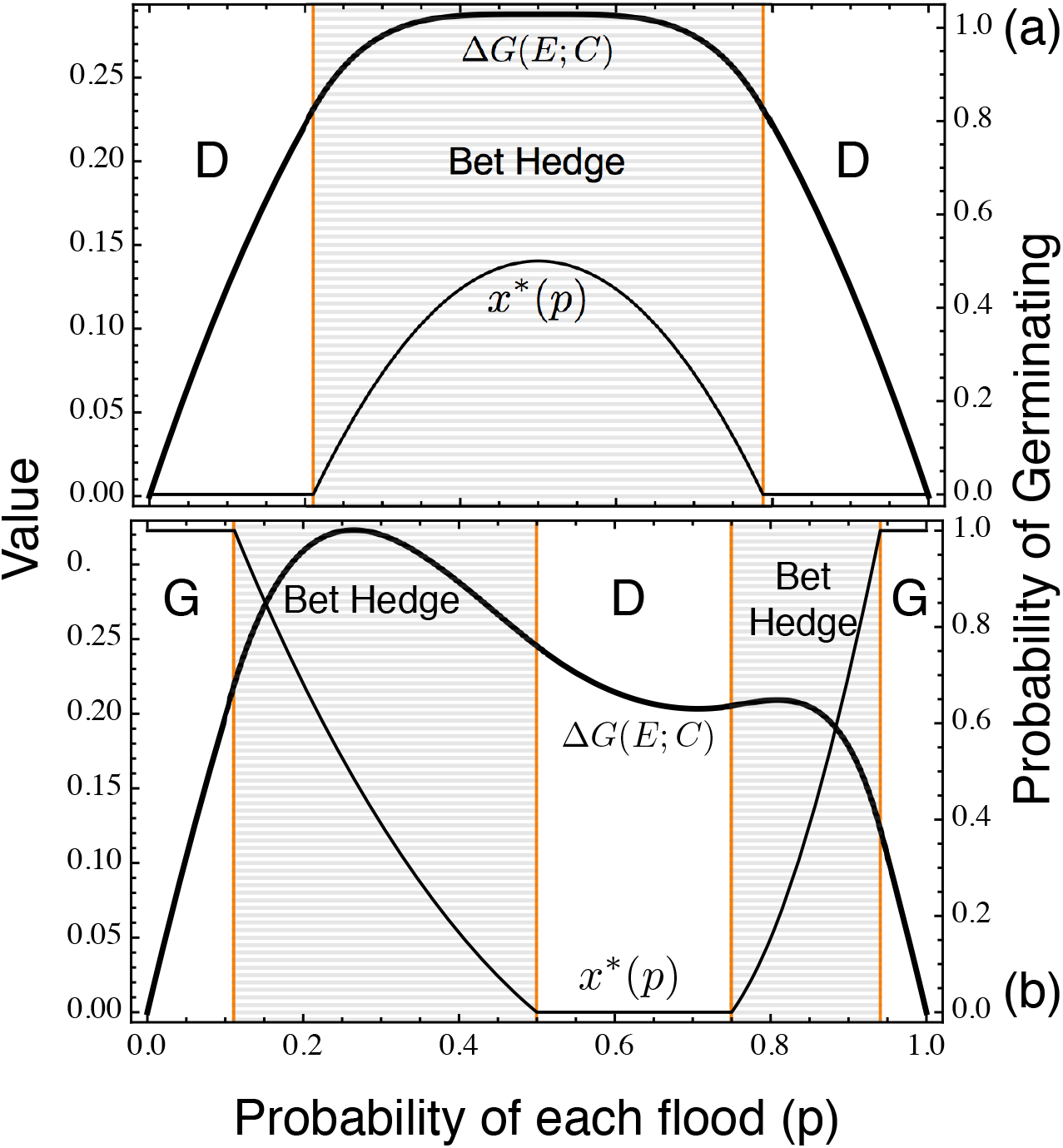
The value of the cue and the probability of germinating as a function of the probability of a single flash flood *p*. Panel (a) is for the example where the intermediate environmental state is favourable for germination. In this case the value of the cue peaks in the region of bet-hedging. The probability of germinating (*x*^∗^(*p*)) increases in frequency but just explores the intermediate regime without ever reaching pure Germination. Panel (b) explores the case where germinating in extreme environments is favourable. This case results in two regions of bet-hedging where again the value of the cue (∆*G*(*E*; *C*)) peaks locally. The probability of germinating goes non-linearly from 0 (Always Dormant) all the way to the other pure phenotype with probability 1 (Always Germinating) hedging its bets along. It changes again in the second region of bet-hedging decreasing to 0 (Always Dormant) where it remains until *p* reaches 1. For both cases, the bounds of the area of bet-hedging can be calculated analytically (ESM). These bounds delineate the transitions in the effective probability of germinating *x*^∗^(*p*).

#### 2.2.2 Adaptation to extremes: Multiplicity of bet-hedging

Contrary to the previous example, it might also be possible that some plants do better in extremes rather than in common environmental conditions. For example, annual pioneer plants can be easily outcompeted by others in intermediate environments (*e*_2_). However at environmental extremes (*e*_1_ and *e*_2_) their seeds have an advantage.

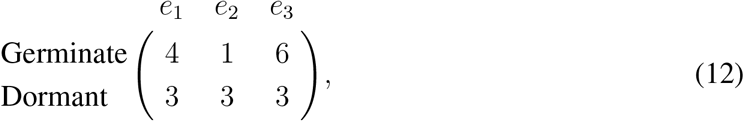

In such a case, we see that the probability of germinating decreases after a specific threshold value of *p* Fig. 3 (b). The non-linear decrease is up to the pure strategy of dormancy. After a specific threshold value of *p*, it is better to hedge bets with a non-linear increasing probability of germination reaching the ‘Always Germinate’ extreme. The value of the cue peaks locally in the bet-hedging regions (Appendix A and Fig. 3 (b) bold curve).

### 2.3 Environmental gradients in the simplex

The probability distribution considered in the previous examples traces a quadratic binomial cline. Such parabolic clines are typical of those distributions in which *e*_2_ is an intermediate environmental state between the others. Thus the distribution of the environments has a direct impact on the bet-hedging regions. For skewed non-linear distributions, it is possible to have multiple regions of bet-hedging. We can extend this case to an arbitrary probability distribution of the environmental states considering three states that occur with probability 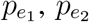 and 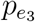, where 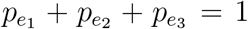. Hence, bet-hedging can occur in a simplex defined by these three probabilities, as shown in Figure 4 (a).

**Figure 4:**
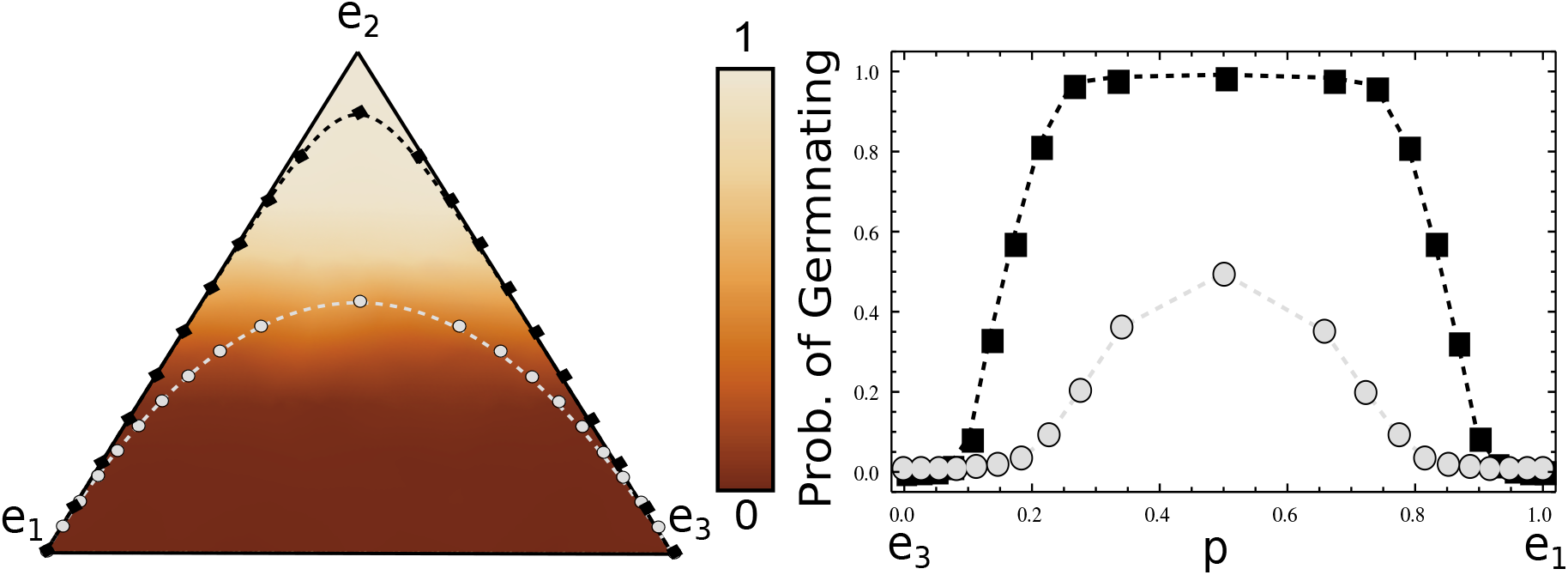
Stochastic simulations of the mean probability of germinating as a function of the distribution of environmental states. Wright-Fisher simulations were performed with a population size, *N* of 1000. After *N* generations the mean probability of germination was calculated. The results were then interpolated to generate the density ternary plot (a). The marks along the clines represent the probability distributions in the simplex space used for the simulations in (b), shown along the clines. Grey dots indicate the second order binomial clines also used for the examples. Black squares indicates instead a fourth order binomial, where the mixed term are assigned to the intermediate state *e*_2_.

We explore the evolution of bet-hedging in the simplex of environmental probabilities by using stochastic simulations for finite populations relaxing the assumption of unbounded exponential growth used for the geometric mean approach. We consider the interaction matrix Eq. (11) and performed Wright-Fisher simulations considering a population size of 1000 individuals, evolved for 1000 generations. Analyzing 100 such realizations in Fig. 4 (a) we show the probability of germination for general environmental distributions. The gradient of the probability of germination was obtained by interpolating a grid of 10000 points for the whole simplex.

In Fig. 4 (right panel) we show further simulations performed for the points marked on the clines in Fig. 4 (simplex). The clines represent the particular probability distributions for the environmental states. For instance, the previous examples considered a second-order binomial indicated by the grey dashed cline, where *e*_1_ occurs with probability *p*^2^ and *e*_3_ with (1 − *p*)^2^. The second black cline indicates the environmental states distributed according to a fourth-order binomial. Thus *e*_1_ occurs with probability *p*^4^ while *e*_3_ with probability (1 − *p*)^4^. The state *e*_2_ occurs with the complementary probability. Depending on the eccentricity of the clines, we can get one or two regions of bet-hedging. We see that the hump shape of the probability of germinating, as predicted by the infinite population size case in Fig. 3 (a), is also recovered via stochastic simulations. For the higher-order cline, the simulation results in the ‘Germinating’ being almost fixed, as is the predicted case on the eccentricity of the cline (Fig. 4 (a)). Hence, the interaction matrix and the particular distribution of the environments influence the observed patterns of bet-hedging.

We explored parabolic clines and represented environmental gradient on a single axis for the sake of simplicity. However, exploring the probabilities of different environments independently in the simplex can be relevant for more complex environments or when different events interact with each other. We provide some examples in an interactive R notebook on Github and Appendix B.

### 2.4 The fitness value of information in presence of constraints

In all previous examples, the fitness value of a cue, ∆*G*(*E, C*), is calculated with an information theory approach, despite the non-linear interactions between environment and phenotypes. Donaldson-Matasci et al. [22] have shown that in the symmetric case, the value is bound by the mutual information between the distribution of an informative cue *C* and that of the environment *E*, *I*(*E*; *C*). Self-information, denoted as *I*(*E*; *E*), is equal to *H*(*E*). Hence, for an entirely informative cue, the fitness value is bound by the entropy *H*(*E*). Does this boundary always hold even in the asymmetric case? If yes, is it possible to draw a lower boundary? Let us consider environmental state *e_i_* with *i* in 1 … *n* and phenotypes *φ_j_* with *j* = 1, … , *m*. Previous models [20, 22] explored cases in which the optimal response to each different environmental state *e_i_* is a phenotype *φ_i_*. In this case, a square *n* x *n* interaction matrix defines the payoff for each phenotype *φ_i_* in each different environmental state *e_i_* [22], occurring with probability *p_i_*. The optimal matches between the environmental state and phenotype appear along the diagonal, the corresponding gains being *a_i,i_* for every *i* = {1, 2, … *n*}:

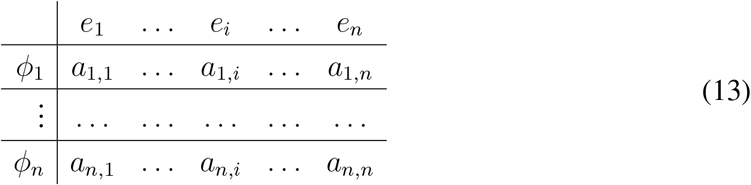

However, the assumption of an optimal, accessible phenotype for each environmental state might not hold, particularly in the context of the evolution to multiple environments, where a single phenotype might represent the best option in response to multiple environmental states. Here we extend the information-theoretic approach developed in Bergstrom et al. [20] to estimate the fitness value of a cue but applied to such cases.

To do this, we start by the proportional betting case, in which the best phenotype response for environmental state *e_i_* provides a payoff *a_i,i_* = *a_i_* while all the other phenotypes are fatal (*a_i,j_* = 0 for *i* ≠ *j*)[22], resulting in a diagonal payoff matrix. In general, the best bet-hedging strategy x, employing a phenotype *φ_j_* with probability *x_j_*, can be determined by solving the Lagrangian [26] of the long term growth rate with constraint (Σ_*i*_ *x*_*i*_ = 1):

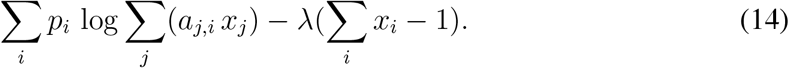

In the proportional betting case, each phenotype is adopted with a probability equal to the occurrence of the environmental state where the payoff is non-zero, denoted as *a_i_*. Hence, in the symmetric case, when for each environmental state a different optimal phenotype exists:

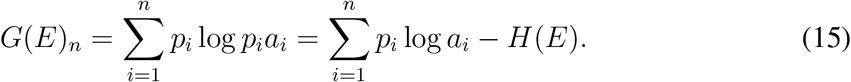

We study the asymmetric case with the number of environments *n* larger than the number of phenotypes *m*. To this aim, we can extend a diagonal payoff matrix *m* x *m* to include additional columns representing the payoffs of the different phenotypes to environments *j > m*. Hence, we investigate payoff matrices for which *a_i,j_* ≠ 0 only when *i* = *j* or for *a_m,j_* and *j* ≥ *m*. We write *a_i,i_* = *a_i_* for *i* < *m*, and *a_m,i_* = *b_i_* for *i* ≥ *m*. We can the write:

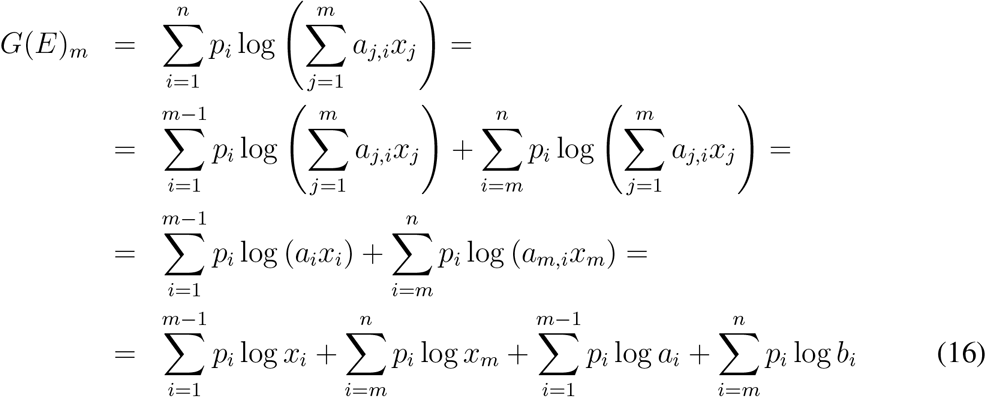

Which gives a maximum at

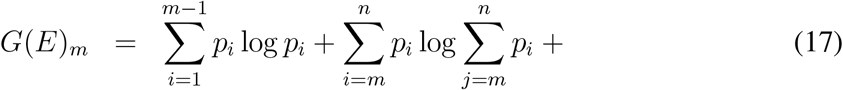

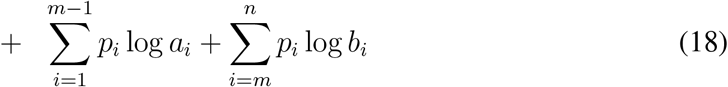

Therefore:

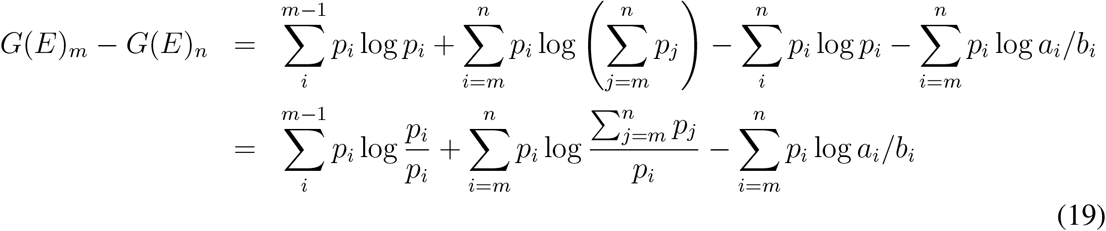

When the cue *C* provides full information about the environment, the right phenotype will be chosen in each case, with payoff *a_i_*, except for environmental states *i > m*, when one strategy cannot develop phenotype *φ_i_*. Therefore just as in the symmetric diagonal case, 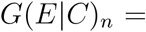 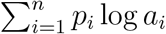, while 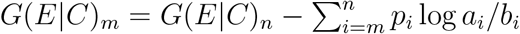 and

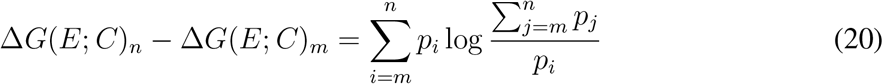

This is the Kulback-Leibler divergence between the optimal strategy when all phenotypes 1 *… n* are available and used with frequency *p_i_*, and when no optimal specific types for environments *m … n* are available, and for all those phenotype *m* is used with the sum frequency 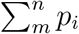. The expression is independent of the payoffs and always positive, confirming that the mutual information bound for the fitness value of an informative cue exists for the asymmetric case. Furthermore, this quantity is part of the conditional entropy of the environment given a fully informative cue, reflecting the residual uncertainty of a cue unable to discriminate between environmental states for which a single phenotype is optimal. Since *I*(*E_n_*|*C*) − *H*(*E*|*C_i_*_≥*m*_) = *I*(*E_m_*|*C*), this indicates that in the presence of a generalist phenotype, the fitness of an informative cue is further decreased by the amount of information necessary to differentiate between the equally paying environmental states. The decrease is equal to the mutual information of the environment with a cue informative only on environmental states for which different phenotypes are optimal.

The above results, shown for the proportional bet-hedging case for simplicity, are also valid in the general case where the payoff of the least advantageous phenotype-environmental state combinations may provide non-zero payoffs. Following Donaldson-Matasci et al., 2008 [38], the fitness profile of these suboptimal phenotypes in different environments can be represented as a mixture of different specialized phenotypes with zero-payoffs for disadvantageous phenotype-environmental state combinations, i.e. as in the proportional bet-hedging case. Briefly, this can be achieved by decomposing a square interaction matrix into the product of two matrices: a diagonal matrix describing the interaction matrix of the proportional bet-hedging case and a stochastic matrix. The latter maps a strategy vector for a general interaction matrix into a strategy vector for a proportional bet-hedging interaction matrix providing the same fitness [38]. To achieve this operation, we need to add rows *m … n* to extend our original rectangular interaction matrix into a suitable and invertible square matrix. However, this is trivial and always possible since the strategy vectors we are interested in have only 0 entries for *i > m*. Thus, also bet-hedging strategies employing non-completely specialized phenotypes can be seen in turn as mixtures of completely specialized phenotypes, allowing us to generalize our results to any interaction matrix. Hence, even in the more complex cases, we see that the cue’s value is bound by the mutual information between the cue and the environment (Fig. 5 and Appendix A). Since we are dealing with perfect cues, this mutual information is simply the entropy of the environment surface) in the asymmetric case). 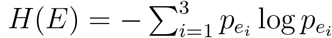 (equal to *H*(*E_m_*) (yellow surface) in the asymmetric case).

**Figure 5:**
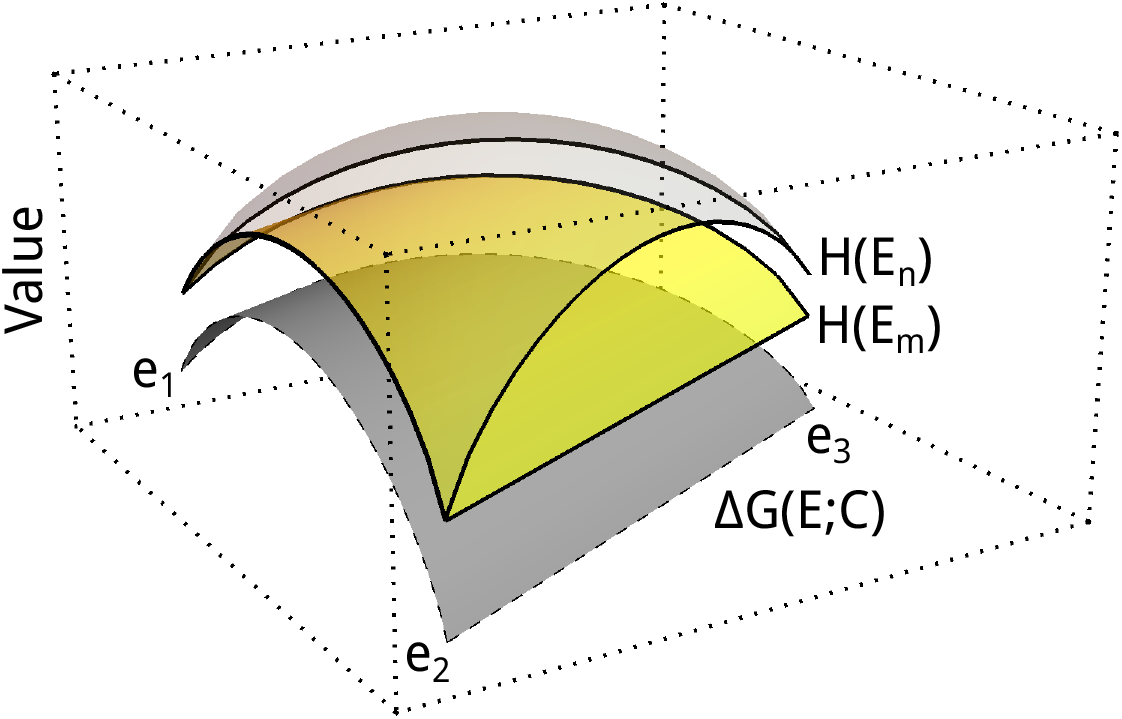
Boundedness of the value of information by the entropy of the environment. Considering all possible distributions, the parameter space of the possible environmental states is then defined on a simplex given by *p*_1_, *p*_2_ and *p*_3_ where the vertices are the pure environmental states *e*_1_, *e*_2_ and *e*_3_ respectively. The value of the cue ∆*G*(*E*; *C*) over such a space (gray triangle) is bounded by the mutual information between the environment and the cue (transparent triangle), that in the asymmetric case decreases to *H*(*E_m_*) (yellow surface). We demonstrate this for the most complicated case described by the interaction matrix 12.

### 2.5 Limits of bet-hedging in increasingly complex environments

So far, we focused on cases in which the number of potential phenotypes *m* is lower than that of environmental states *n*, i.e. payoffs matrices with *m* < *n*. We now describe why these cases are relevant and likely universal in nature by showing that even when more potential phenotypes are available and *m > n*, evolution leads to organisms using only a reduced set of phenotypes. Note that non-trivial instances of *m > n* matrices correspond to cases in which at least one phenotype provides a non-zero payoff for multiple environmental states, thus offering an opportunity for conservative bet-hedging. An extreme case is that of a phenotype that provides the same payoff for all environmental states. An alternative case is of an intermediate phenotype that can be adopted despite two or more optimal phenotypes for specific environmental s tates. Hence, our question corresponds to investigating when conservative bet-hedging strategies evolve over diversifying bet-hedging ones, i.e. when a matrix *m × n* reduces effectively to a matrix with a smaller number of rows/phenotypes. To do this, we examine a matrix:

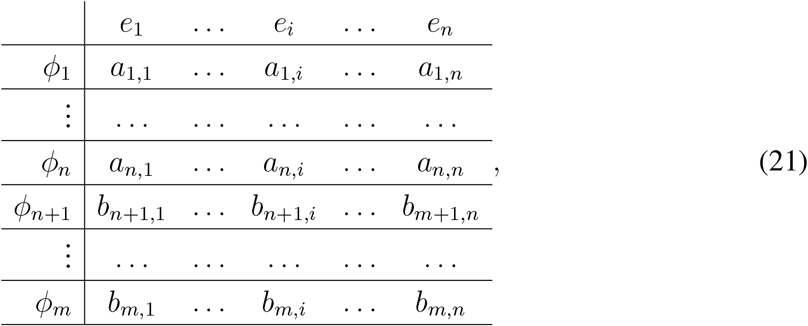

where for each environmental state *i* we arranged the phenotype providing the highest payoff as phenotype *j* = *i*, with payoff *a_i,i_*. Further potential phenotypes are denoted as *φ_j_* with *j > n*, and by construction, payoffs *b_j,i_ ≤ a_i,i_*. Our previous analysis presents a simple method for investigating under which environmental circumstances such asymmetries might evolve. Consider a strategy *G*(*E*)_*m*_ adopting a generalist phenotype *φ_j_* with payoffs *b_j,k_ > a_i,k_* rather than the specialized phenotypes *φ_k_* with payoffs *a_k,k_* for *k > i*, as strategy *G*(*E*)_*n*_. By recalling Eq. (19)-20, we can see that *G*(*E*)_*m*_ is advantageous if

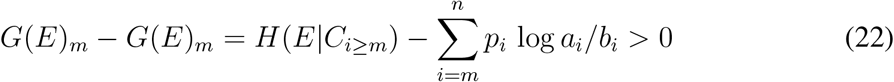

indicating that the maximum decrease in payoff of the generalist strategy is bound by the entropy of the system that is *removed* by adopting it. This simple relationship helps us to illustrate the instances in which generalist phenotypes are employed. To do this, we devise an example in which specialized, intermediate and generalist phenotypes evolve, following the general payoff matrix described in Eq. (21). We consider *n* equally frequent environmental states, subdivided in *n_k_* groups of size *k*, biologically representing environmental states with similar features. Intermediate phenotypes allow to respond to any of the similar environmental states, but with maximum payoff lower than the specialized phenotypes, i.e. for each of the *n_k_* groups, an intermediate phenotype *φ_i_*_≥*n*_ exists, providing payoff 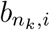 as in Eq. (21). Also, a perfect generalist phenotype exists, providing the same payoff *c* in all environmental states.

By manipulating the number of environmental states, while keeping constant the size of the groups, the total entropy (*H*(*E_n_*)) of the system increases, while the entropy within the groups (*H*(*E*(_*k*_)) is not affected (Fig.6a). Furthermore, the conditional entropy of the environmental states with the same intermediate phenotype is unaffected; intermediate phenotypes are not favoured over specialized ones as the total entropy increases. Whether intermediate phenotypes evolve or not depends exclusively on the ratio of the payoffs. On the other hand, generalist phenotypes (i.e. conservative bet-hedging strategies) eventually become advantageous as entropy increases.

**Figure 6:**
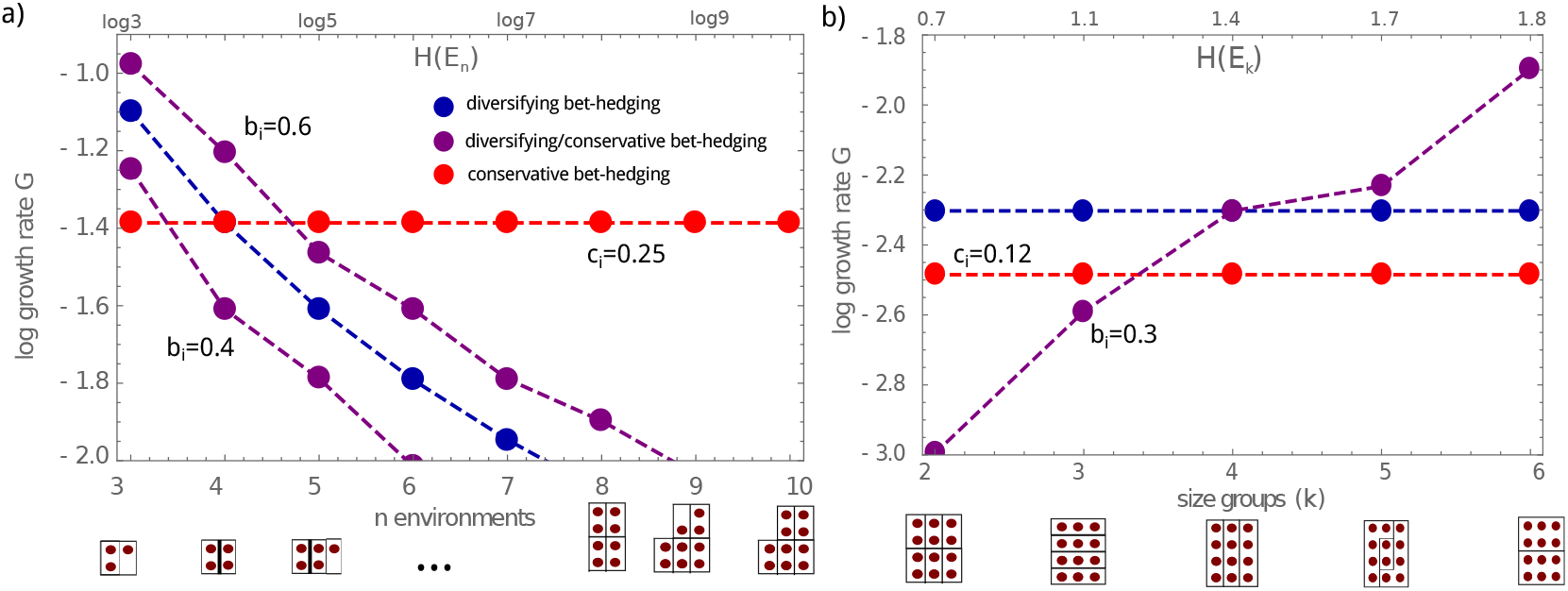
Limits of bet-hedging in increasingly complex environments. Long term log-growth rates for different bet-hedging strategies for (a) environments with an increasing number of uniformly distributed environmental states or (b) an increasing size of the partitions of environmental states for which an intermediate generalist phenotype exist. These partitions are represented under the plots with environmental states (brown circles) grouped based on conceivably relevant generalist phenotype (represented as black rectangles). We assume a square environment payoff matrix in which the optimal payoff for any environmental state give payoff 1 (*a_i,i_* = 1) and 0 in all non-matching cases (*a_i_*_≠*j*_ = 0), leading to a fully diversifying bet-hedging strategy (blue dots). We also show the log-growth rate for a fully generalist strategy with a fixed payoff in all environmental states *c* (red dots) and partially bet-hedging strategies that instead of hedging their bets exclusively on the optimal phenotypes, they hedge more generalist phenotypes matching the different partitions, providing payoff *b* in those and 0 in the rest. a) Partition size is fixed to two.

By manipulating the size of the groups, only the conditional entropy, *H*(*E*|*C_i_*_≥*m*_) (here *H*(*E_k_*)), of the groups increases (Fig.6b). In this case, the growth rates of pure diversifying and conservative bet-hedging remain constant. However, strategies hedging their bets between intermediate phenotypes become more advantageous as *H*(*E_k_*) increases.

To summarize, in both cases, strategies employing generalist phenotypes evolve as the uncertainty of the system increases. In turn, the growth rate and the fitness value of information can be described using asymmetric matrices.

## 3Discussion

Organisms facing complex and variable environments often evolve complex adaptive mechanisms to acquire information about the environment from informative cues. What is the fitness values of these cues? What is the fitness value of these complex adaptations? Answers to these vital questions remain elusive in natural populations. Information theory is a useful framework to answer these questions. First, it provides a theoretical boundary and an upper estimate of the fitness value of cues and phenotypes. Second, our approach helps leverage measures and features of the environment, specifically its uncertainty, and may help ease the design of complex ecological experiments. This knowledge can further inform empirical as well as theoretical research. Despite these advantages, the application of this theory has been so far limited [21], possibly because the models usually focused on simple tractable systems. Natural systems, on the other hand, can be exceedingly complex.

Organisms have evolved sophisticated bet-hedging strategies to deal with environmental uncertainty without relying on informative cues. In these cases, organisms do not need to respond flexibly to the environment. Spreading the risk of facing adverse environmental conditions by betting on diverse or intermediate phenotypes can work. This mechanism can explain several biological phenomena, from intergenerational phenotypic stochastic switching to animal personalities [39], although hard evidence for bet-hedging in the animal world has been so far elusive [14]. While it is possible to study bet-hedging experimentally in microbes [6, 7], the evolutionary mechanisms underpinning the use of informative cues is still elusive. Different levels of environmental unpredictability present along environmental or geographical clines exacerbate the complexity of bet-hedging and informative cues. Natural populations could then potentially display non-trivial bet-hedging patterns. We illustrate this through the example of organisms adapted only to intermediate states of environmental gradients such as temperature or salinity. Intermediate levels of such gradients are taken as discrete environmental states. For example, seasonal plants adapted to intermediate water levels: the total absence of flash floods might lead to a harsh dry year, and two consecutive floods may damage seedlings. In this example (Fig. 3), we show two cases in which bet-hedging occurs only within confined regions of the parameter space, i.e. regions with intermediate probabilities of rain, surrounded by more stable regions where only one phenotype is adopted. These hypothetical clines, described in Figures 4, could be geographic transects going from environmental extremes from desert to rainforests. For example, dormancy is advantageous in response to multiple extreme events (e.g. drought, fire [40], extreme cold) and plants occurring in many different environments, from alpine forests [41] to wetland [42] and deserts. We suggest that such studies could highlight interesting patterns of bet-hedging along environmental gradients, primarily if a species exists across such environmental transect [43].

We show that the fitness benefit of an informative cue is universally bound by the mutual information between the cue and the environment. Besides, when the variety of phenotypes that can be adopted by phenotypic switching strategies and plasticity is limited, the maximum potential fitness benefit of an informative cue is further reduced. Precisely, the reduction is due to the conditional entropy between the environment and a cue informative about the inaccessible phenotypes. This result implies that plasticity is less advantageous when physiological constraints limit the phenotypes that can be adopted. Although intuitive, this observation has intriguing implications: we have seen that as environmental uncertainty increases, the number of adoptable phenotypes by a bet-hedging organism can decrease; then, even if this organism has information to develop one of the available phenotypes, the fitness benefit of the informative cue can be lower, than the benefit to an organism living in a simpler environment. In practice, under high environmental variation, an organism might decrease the amount of explored phenotypes explored through bet-hedging, further reducing the benefit of relying on informative cues.

Note that information theory provides insights into many possible scenarios in which the relationship between bet-hedging and cues is complicated. For example, when cues are unreliable and idiosyncratic (i.e. each individual - a different cue), relying on cues can be used as another means towards bet-hedging [19]. Hence, we advocate that collecting more empirical data on the environmental uncertainty of species that live on environmental gradients might provide exciting insights into bet-hedging. Such studies would allow a quantification of the fitness benefits of informative cues using the framework described above. To this aim, we showed in the examples and the accompanying R notebook (Github and Appendix B) how such data could be analyzed. We emphasize that this approach is general, and in the text we used parabolic environmental clines only as an example. Conversely, the approach applies to cases in which multiple environmental states and environmental factors could affect the fitness of an organism.

For instance, density dependence and competition are essential predictors of bet-hedging [44], as in advancing pioneer plants, in which dormancy can evolve in response to the competition with plants adapted to prevailing environmental conditions. Most importantly, this approach provides theoretical upper limits for the fitness advantage of specific cues or adaptive mechanism even in the absence of precise knowledge of fitness, a task that is otherwise hard to achieve. To illustrate this, we look at a hypothetical case inspired by the marine midge *Clunio marinus* (Chironomidae, Diptera). Clunio is adapted to one of the most complex environments on earth, the intertidal zone of seacoasts. Synchronizing the adult emergence time to multiple environmental cycles of tidal, diurnal and lunar cycles, the marine midge has successfully adapted in processing complex informative cues [45, 46, 47]. Using cues, adults emerge when conditions are favourable. Assuming a narrow window of opportunity for successful reproduction, we provide a theoretical upper boundary for the fitness advantage of this trait equivalent to about 5 bits for the whole life-cycle, about 3 bits due to lunar cues and 1.8 to circadian ones (see Github or Appendix B for additional explanations and examples). Providing a metric in bits, valid across species, a thorough quantification of environmental variability in different species along environmental gradients could allow for a clearer understanding of the ecological conditions promoting plasticity and the usage of informative cues.

In conclusion, we have extended the study of bet-hedging to asymmetric cases where phenotypes are scarcer than the number of different environmental states. We show that in realistic conditions, the patterns of bet-hedging might be more complicated than previously expected. The results could help clarify some of the mixed results of empirical observations [14], and better describe the evolution of bet-hedging along complex environmental gradients. However, even the most complicated cases still obey the limit on the fitness value of a cue imposed by mutual information.

## Acknowledgments

CSG acknowledges Arne Traulsen for helpful discussions and funding from the New Zealand Institute for Advanced Study, Massey University, where part of the work was done. All authors acknowledge funding from the Max Planck Society.

## APPENDIX A Examples with asymmetries in strategies and environments

## Adaptation to intermediate environments

If the difference in benefits in the three environments switch their signs, this satisfies a necessary condition for getting two internal fixed points [48, 49]. Herein, we set up an example of two internal fixed points for the three environment cases. Here

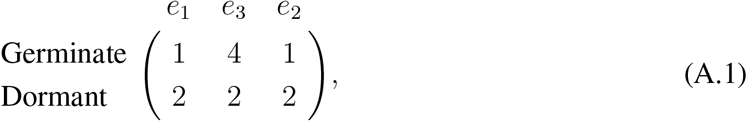

The fitnesses of the two phenotypes are,

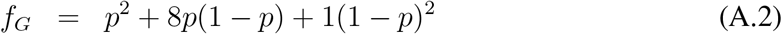

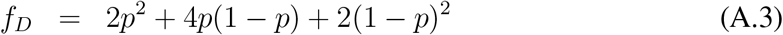

In the absence of any external information about the environment the single generational expected fitness is *F* (*E*) = *max*[*f_G_, f_D_*]. However with a perfectly informative cue *C*, the single generational expected fitness, *F* (*E | C*) = 2*p*^2^ + 8*p*(1 – *p*) + 2(1 – *p*)^2^. We look at the difference between the fitness with perfect information and the strategy of probabilistic allocation according to the game-theoretic outcome. The value of information is measured as the difference between the fitness with the cue and without the cue ∆*F* (*E*; *C*) (Figure A.1).

The frequency with which seeds germinate is assumed to be *x* while they remain dormant with a complementary probability. The average fitnesses of the population in the different environments is then given by,

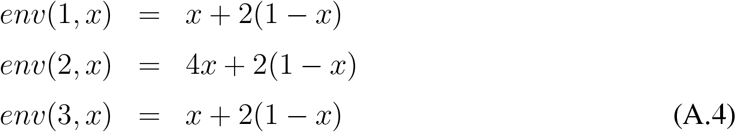

Maximising the long term population growth rate means maximising,

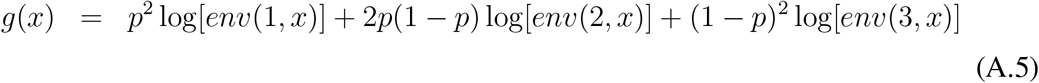

The maxima of this function appears when *x_max_* = −1 + 6*p* − 6*p*^2^. Restricting to values of *p*

**Figure A.1:**
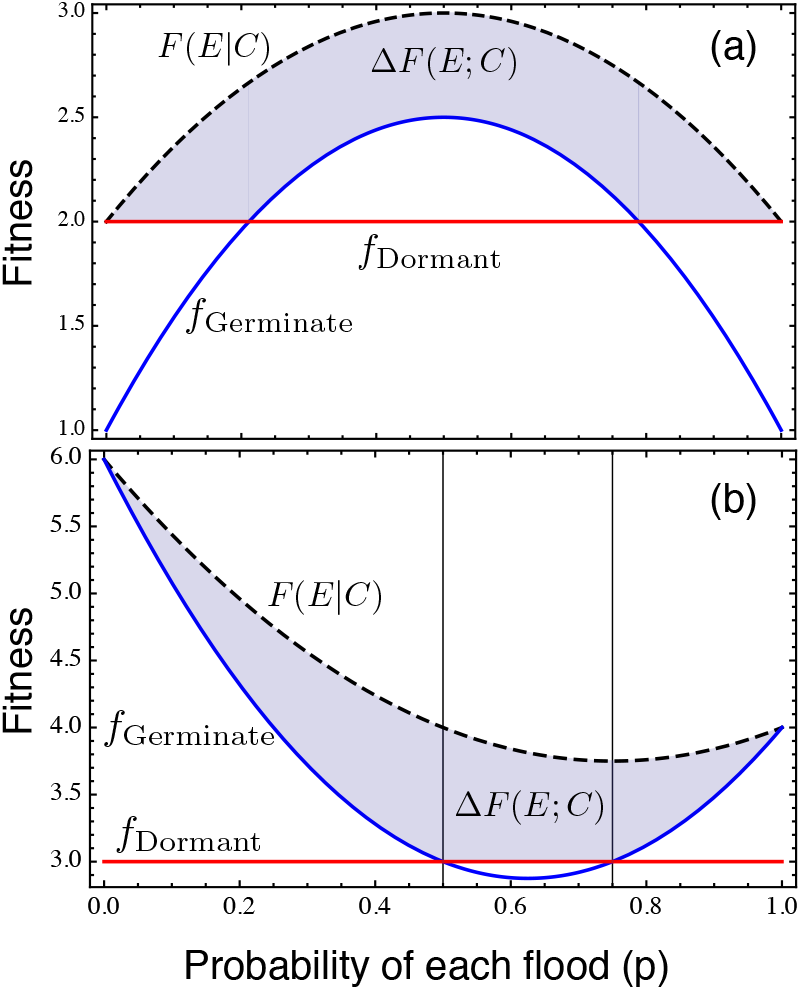
Average fitnesses of the strategies Germinate and Dormant as a function of the probability of a single flood event *p*. Panel (a) describes the situation where the intermediate environmental state is the only one favourable for germination. However knowing the exact state of the environment can lead to an increase in fitness given by *F* (*E*|*C*). Hence the fitness benefit due to the cue, ∆*F* (*E*|*C*), is given by the shaded area. Panel (b) describes the case where the optimal gains for germinating are obtained in extreme environments. For this particular example the we get two internal fixed points given by {1/2, 3/4}. Below and above these fixed points it is best to germinate but in between it pays to stay dormant. Again for a perfect cue the fitness is given by increases by ∆*F* (*E*; *C*) to *F* (*E*|*C*).

which give us results for *x* between 0 and 1 we get,

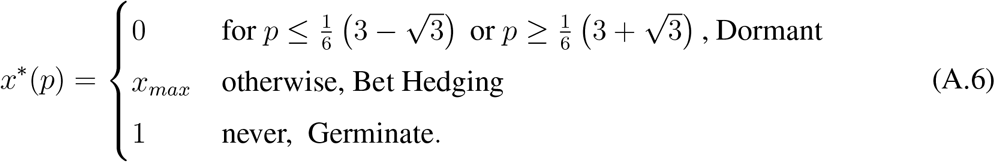

The growth rate without the cue therefore is given by,

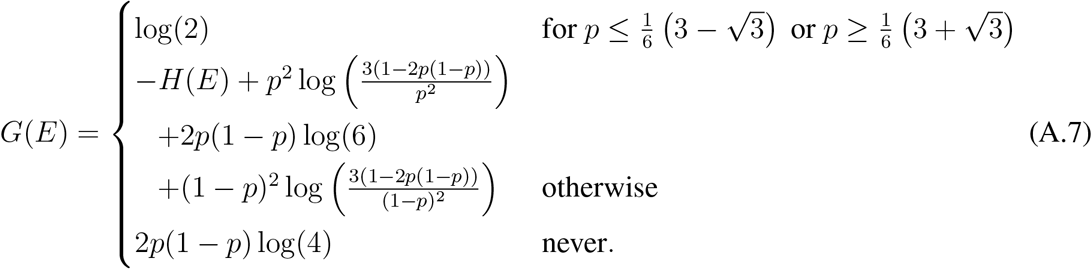

If the cue provides perfect information then the long term fitness value is, *G*(*E*|*C*) = *p*^2^ log(2)+ 2*p*(1 − *p*) log(4) + (1 − *p*)^2^ log(2). The gain in fitness due to knowing the environment, i.e. the fitness value of information is therefore given by, ∆*G*(*E*; *C*) = *G*(*E*|*C*) − *G*(*E*),

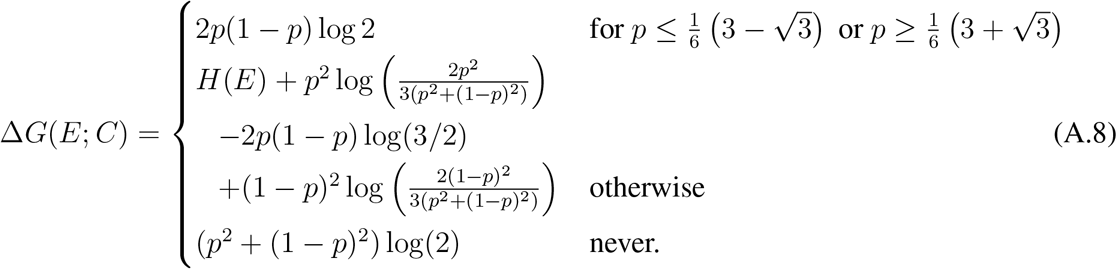

The bet-hedging region in this particular case explores the mixed phenotype space between the pure phenotypes. Therefore in the bet-hedging region, such “exploratory bet-hedging” occurs that would never be affordable as a pure strategy.

## Adaptation to extremes: Multiplicity of bet-hedging

Contrary to the previous example, now we consider seeds that perform better germinating in extreme environments.

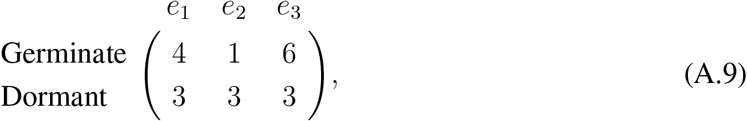

Equating the average fitness of the two phenotypes which are given by,

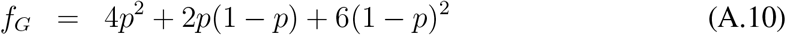

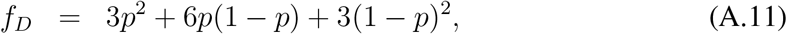

we get two fixed points denoted by *p*^∗^ = {1/2, 3/4}. Thus depending on *p*, the best phenotype, according to the fitnesses, switches from germination to dormancy to germination again.

However if the cue provides perfect information about the environment then the fitness conditioned upon the cue is *F* (*E | C*) = 4*p*^2^ + 6*p*(1 – *p*) + 6(1 – *p*)^2^. The value of the information received due to the cue is the difference between the fitness with the cue and without the cue ∆*F* (*E*; *C*) (Figure A.1). Assuming that the seeds germinate with probability *x* and stay dormant with probability 1 – *x*, the average fitnesses of the population in the different environments are then given by,

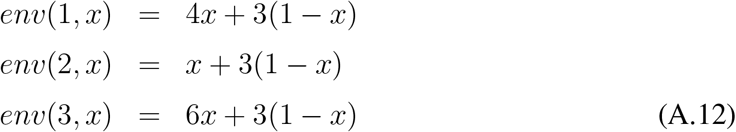

Maximising the long term population growth rate, we get

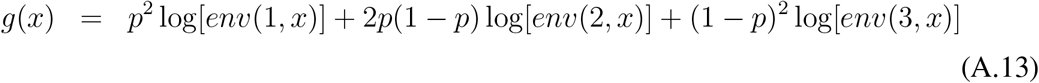

whose maxima is given by,
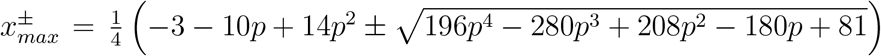. While one of the solutions is outside the range of x (0 ≤ *x* ≤ 1) the other results into the following piecewise solution of *x*^∗^,

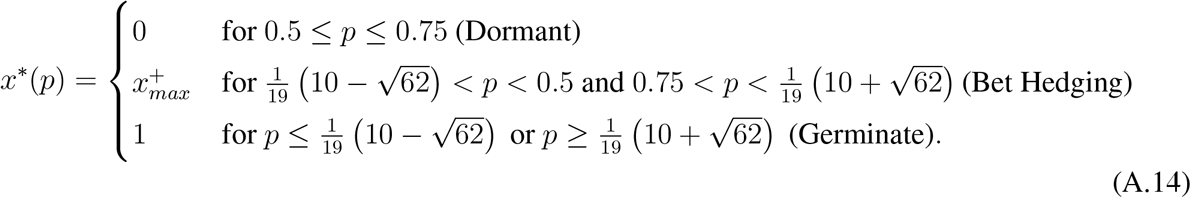

This shows an interesting situation where there are two regions of bet-hedging. The first one changes the phenotype from pure germination to pure dormancy and vice versa in the second region (Fig. A.1). The growth rate without the cue, therefore, is given by,

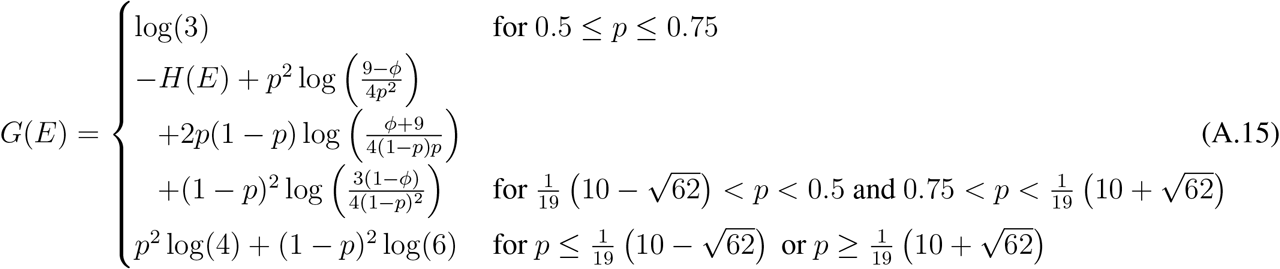

where we use the abbreviation 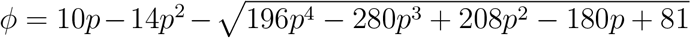. For a perfectly informative cue the long term fitness value is, *G*(*E | C*) = *p*^2^ log 4 + 2*p*(1 – *p*) log 3 + (1 – *p*)^2^ log 6. The gain in fitness due to knowing the environment, i.e. the fitness value of information is therefore given by, ∆*G*(*E*; *C*) = *G*(*E*|*C*) − *G*(*E*),

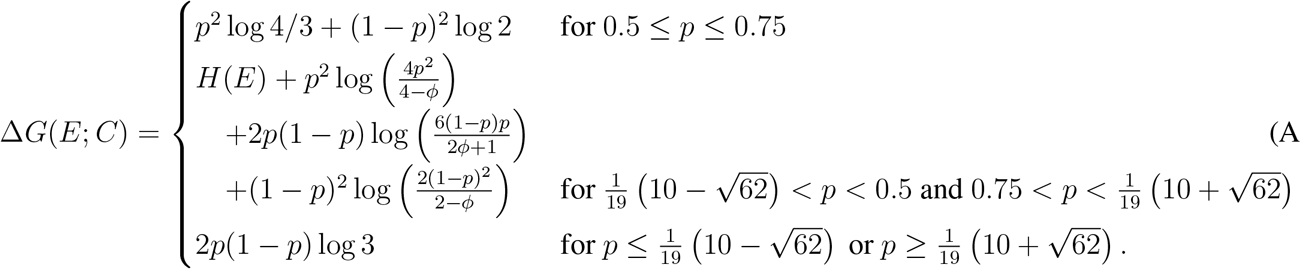

Thus even for this complicated case with two regions of bet-hedging, we can see that the cue’s value is a composite of the mutual information between the environment and the cue and a non-linear function in *p*, which can be interpreted as the probability of a single flood.

## APPENDIX B Examples of estimates of the fitness of informative cues in R code

In this notebook we provide an R tutorial to apply an information theoretic approach to empirical studies from literature. It can be also downloaded in Rmarkdown format from the github repository: https://github.com/tecoevo/informativecues (https://github.com/tecoevo/informativecues)

We use the package “optimx” for numerical optimization. We start by defining useful functions for our estimates:

- G_fx: function to calculate the log growth rate of a bet-hedging strategy adopting a set of phenotypes with probabilities given in the vector pr_ph to respond to a set of environments occurring with probabilities given in the vector pr_e. The payoff matrix A describes the fitness of an individual for each combination phenotype(rows)-environment(columns).
- GC_fx: function to calculate the log growth rate of a strategy when the environmental cues are known, given a set of environments occurring with probabilities given in the vector pr_e and a payoff matrix A. Note that this formula is valid for organisms in which fitness is determined by the occurrence of a single environmental state in a specific time of their life-cycle (as in annual plants). For different cases see the *C/unio* examples below.
- find_optimal_pr.ph: function to estimate optimal bet-hedging strategy by maximizing the geometric fitness. Arguments are a set of environments occurring with probabilities given in the vector pr_e and a payoff matrix A.
- delta_GC_fx: wrapper function that returns the fitness benefit of informative cue G(EIC)-G(E), G(E) and the optimal bet-hedging strategy. Arguments are the same as GC_fx.

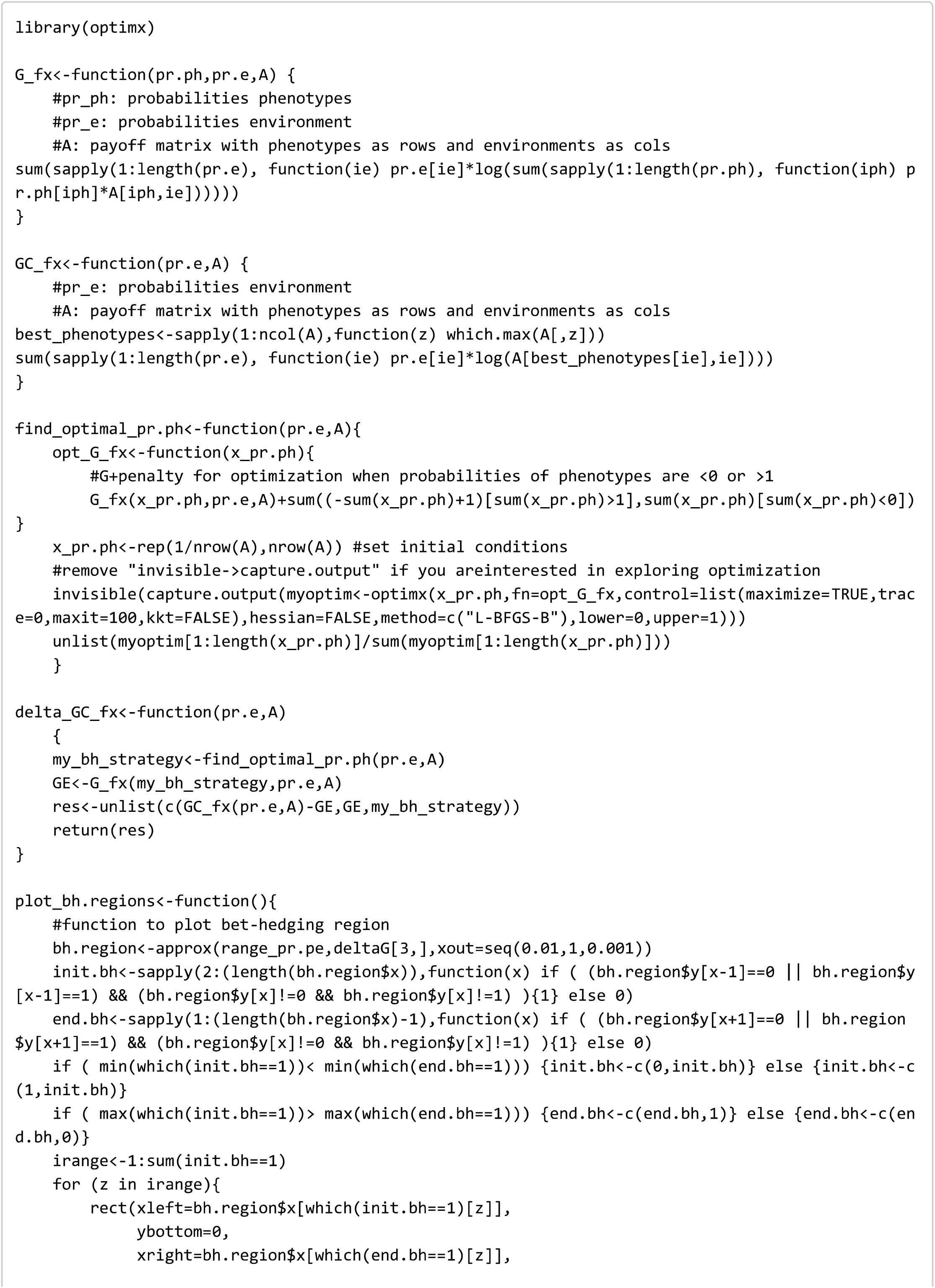

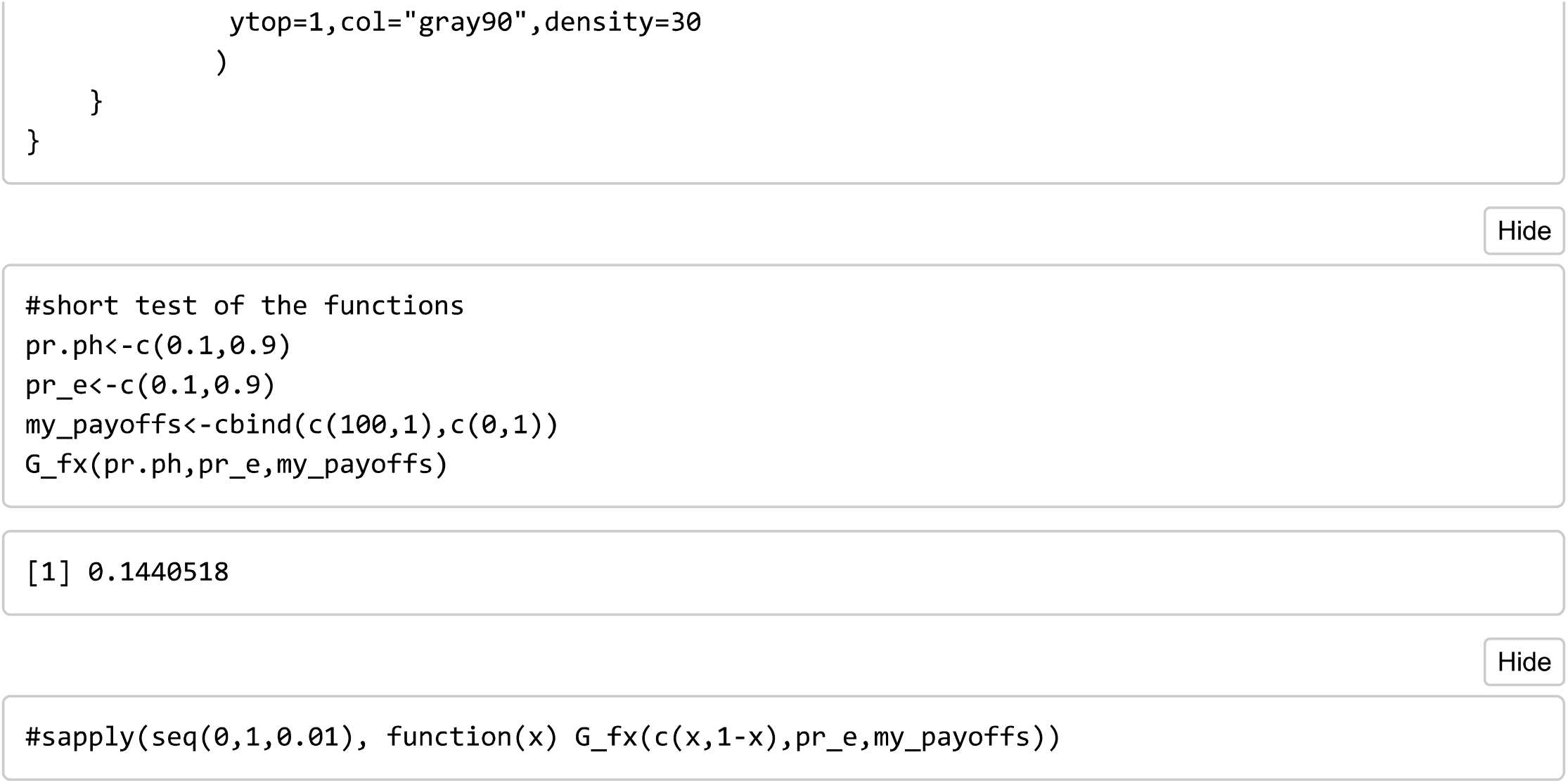

Below we show an example to see that the optimal bet-hedging strategy is not affected by different payoffs in the proportional bet-hedging case with lethal non-optimal phenotypes, while in presence of cues the fitness changes.

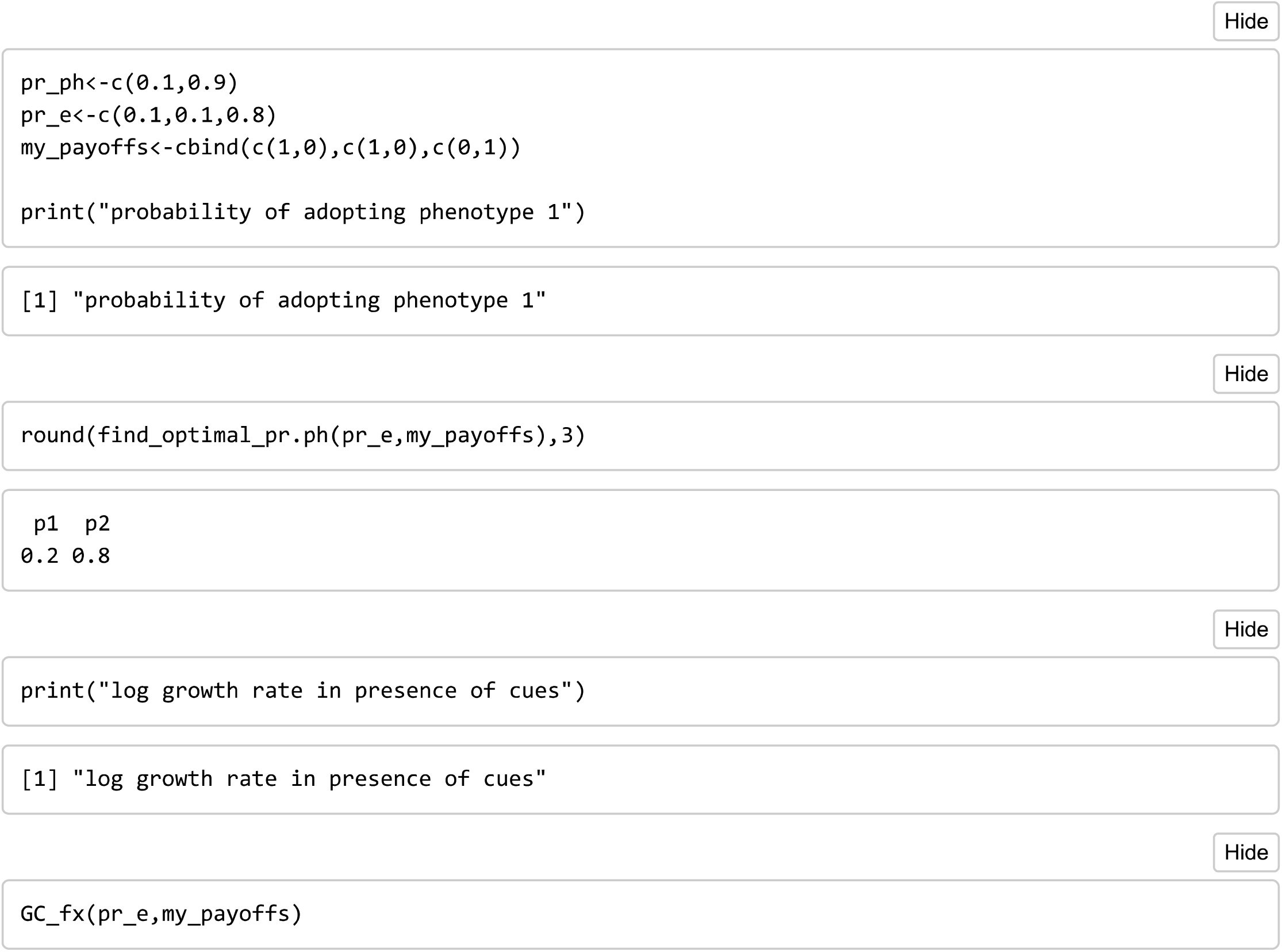

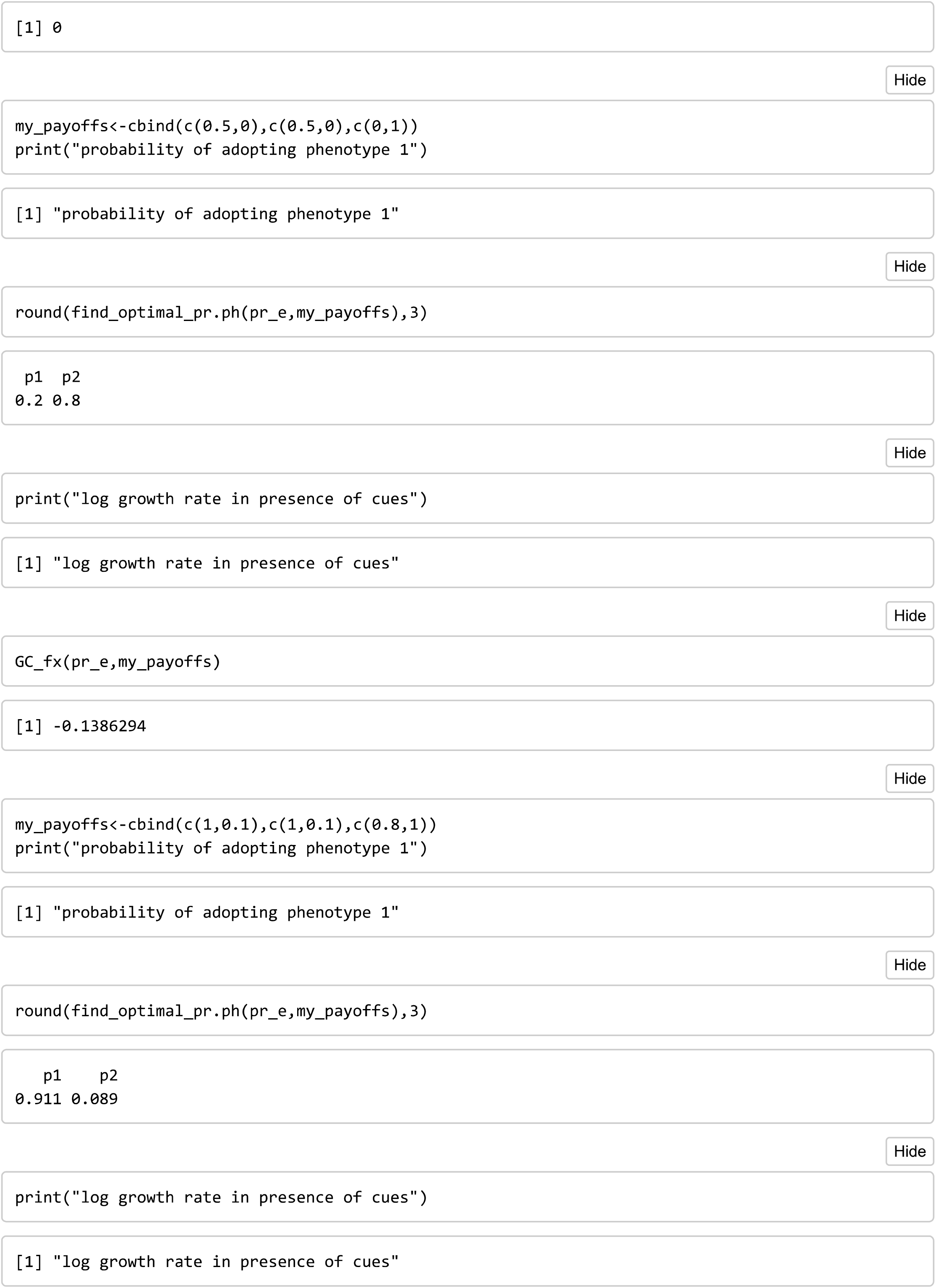

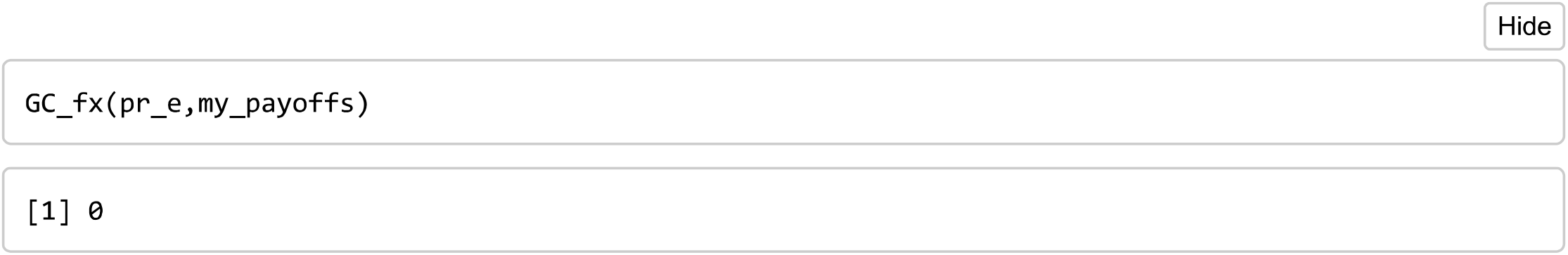

## Floods and desert plants

(Examples from Mafessoni, Lachmann and Gokhale, 2020)

## Symmetric case: two environmental states with two phenotypes

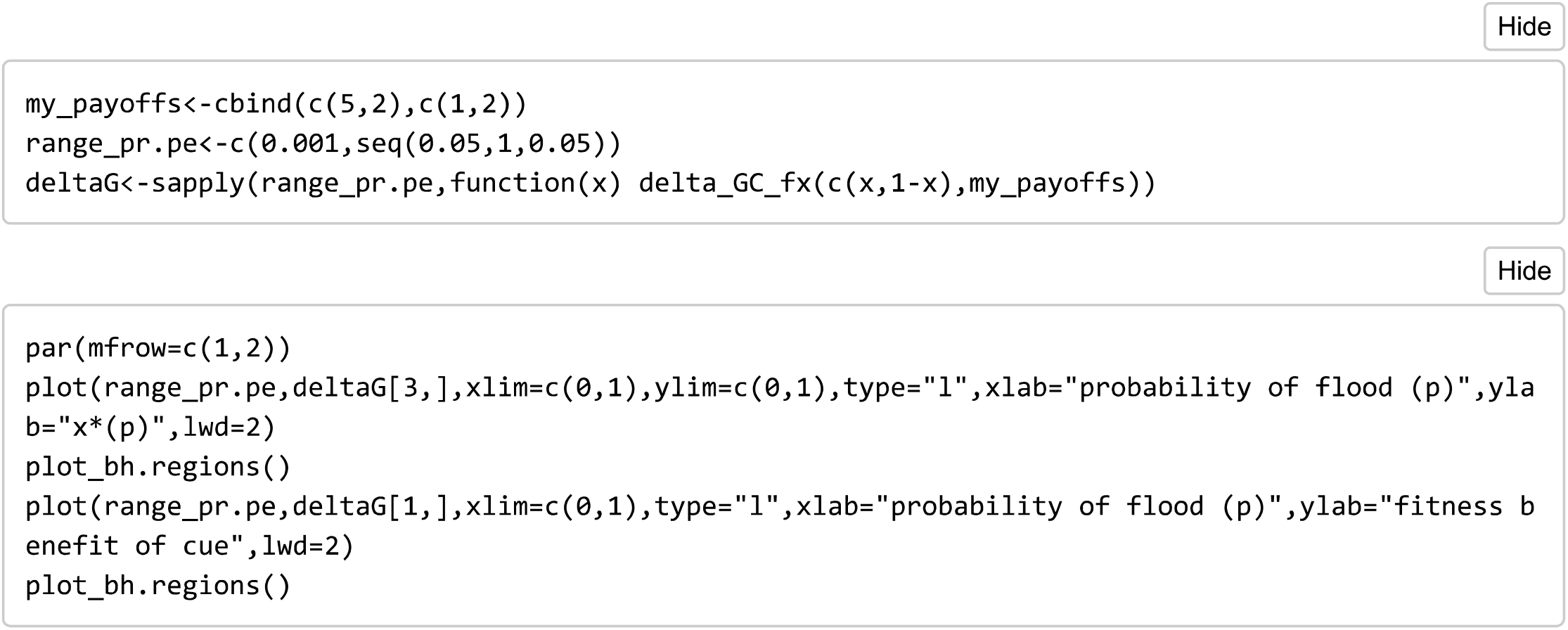

## Asymmetric payoff matrices (3 environments and two phenotypes)

We examined in the main text a classical example of plants exposed to the risk of floods. It is convenient to for them to germinate after a flood, however if a second flood occurs straight after it might wash them away or kill them. In this case we can represent our system along a single probability (p - the risk of floods).

## Adaptation to intermediates

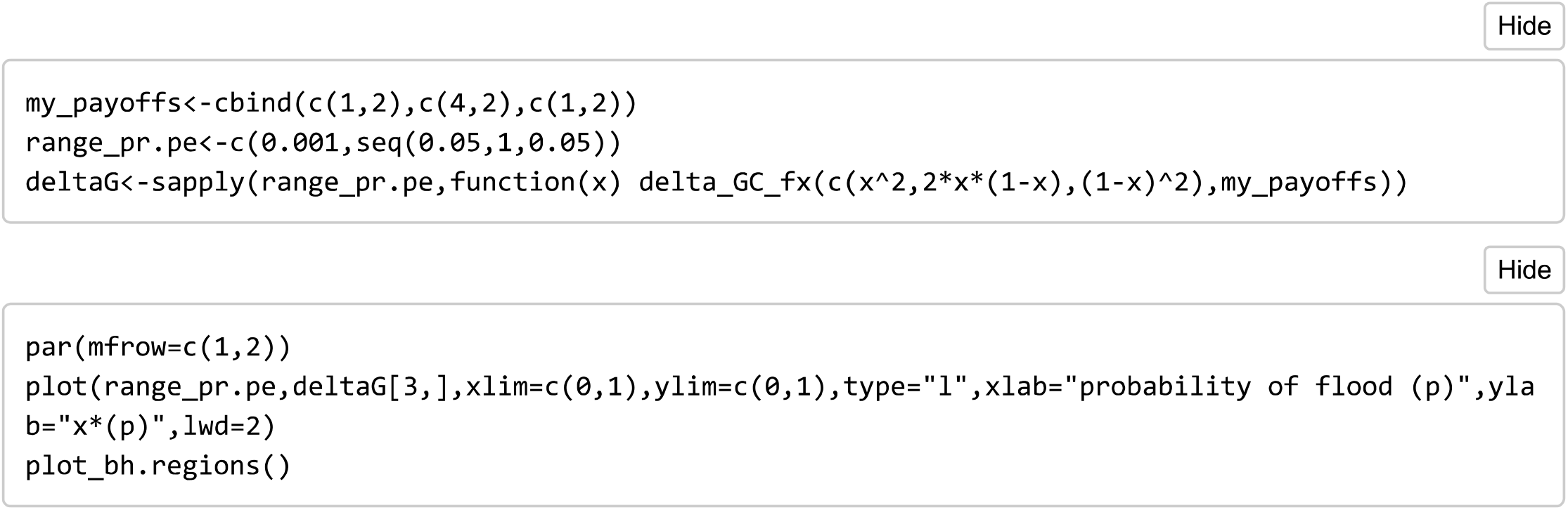

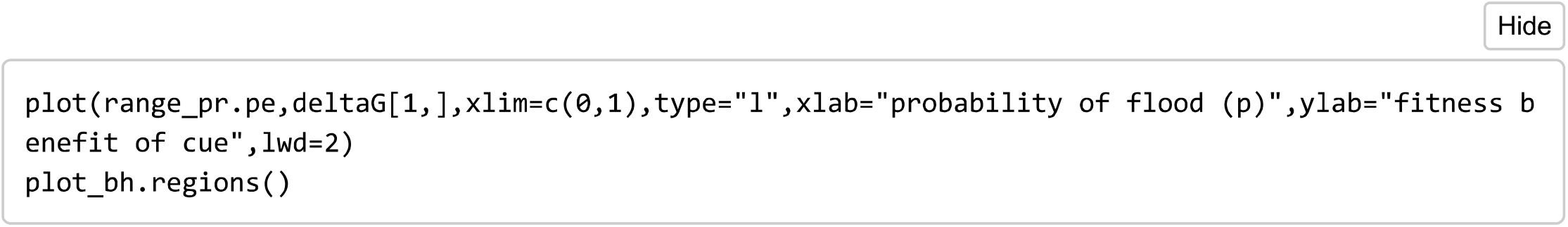

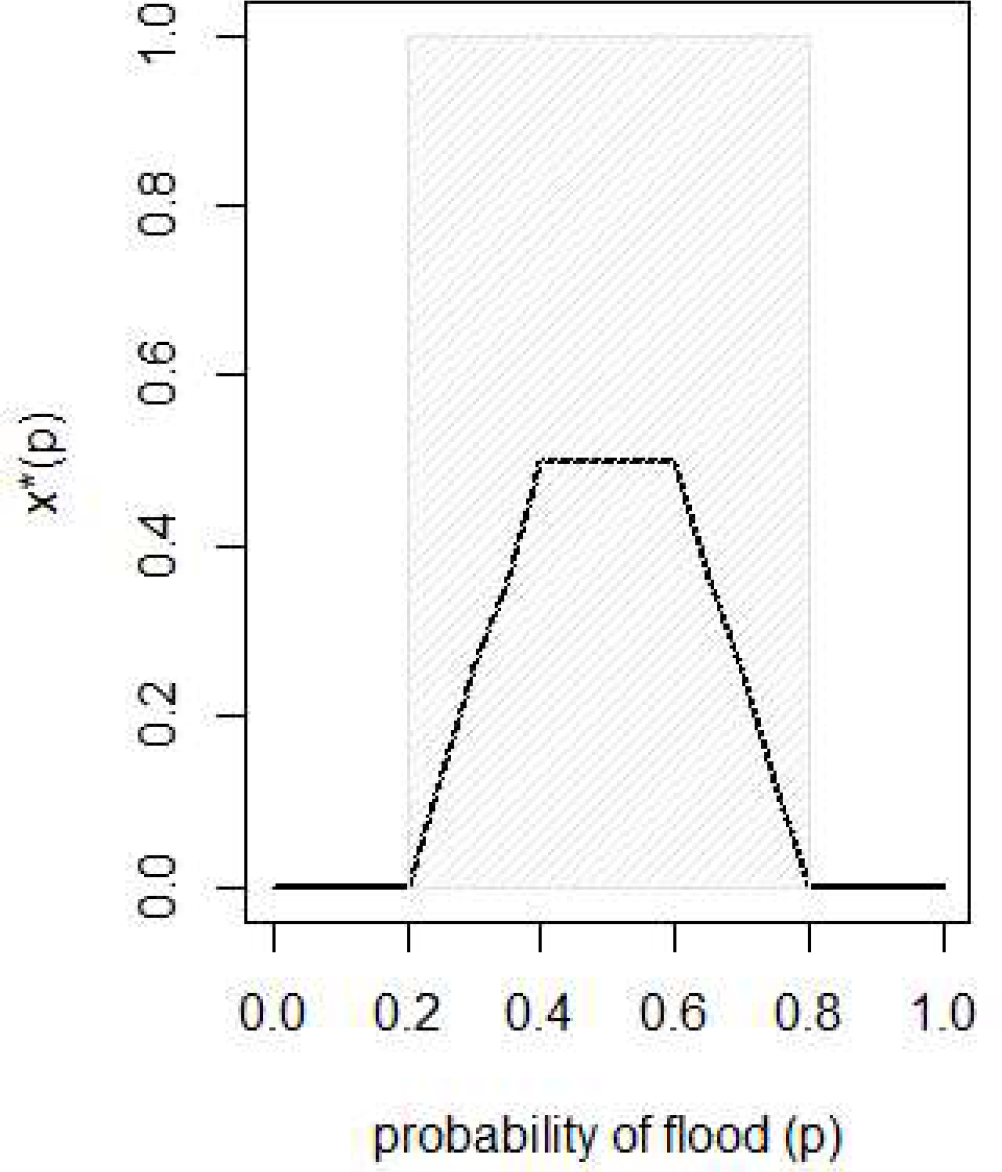

## Adaptation to extremes: Multiplicity of bet-hedging

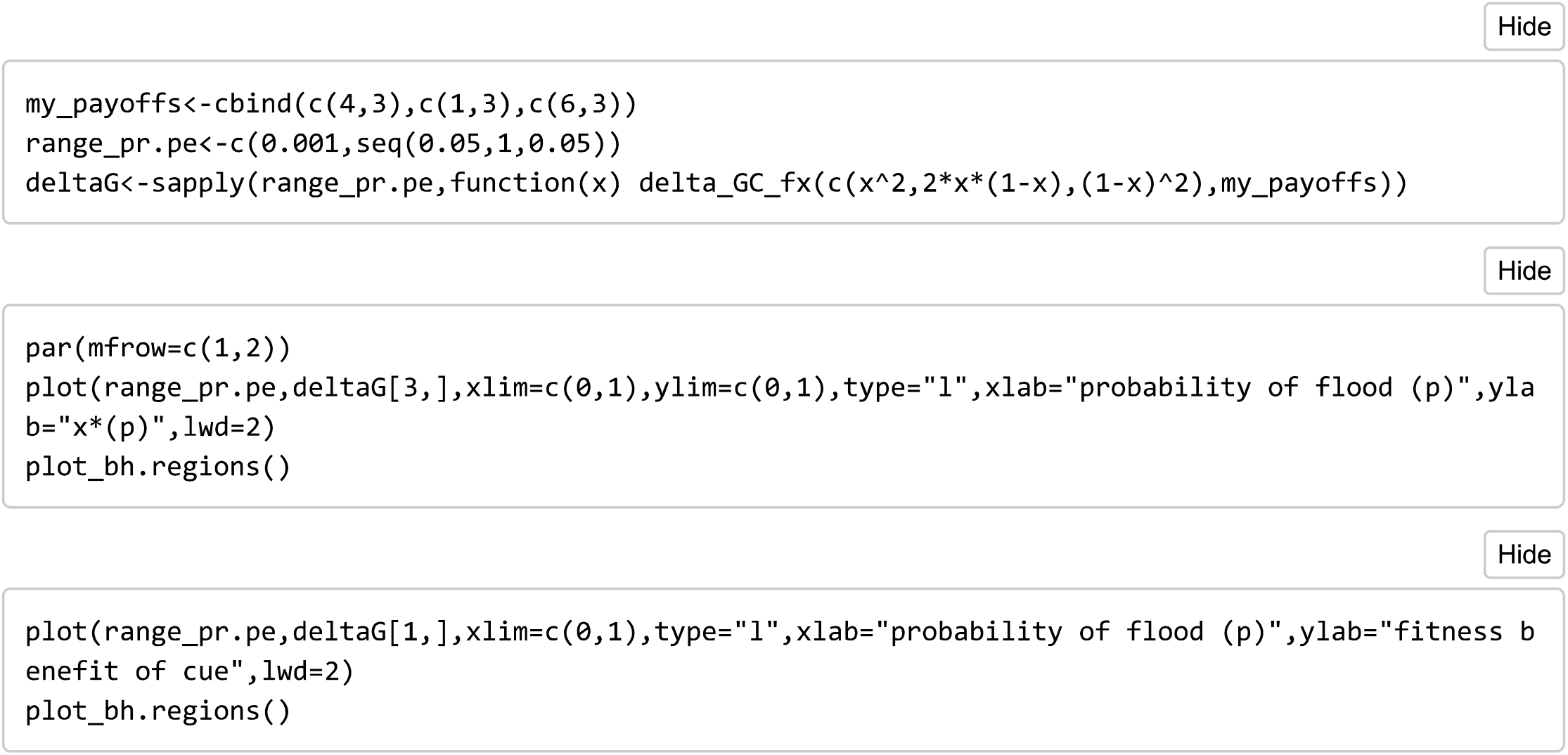

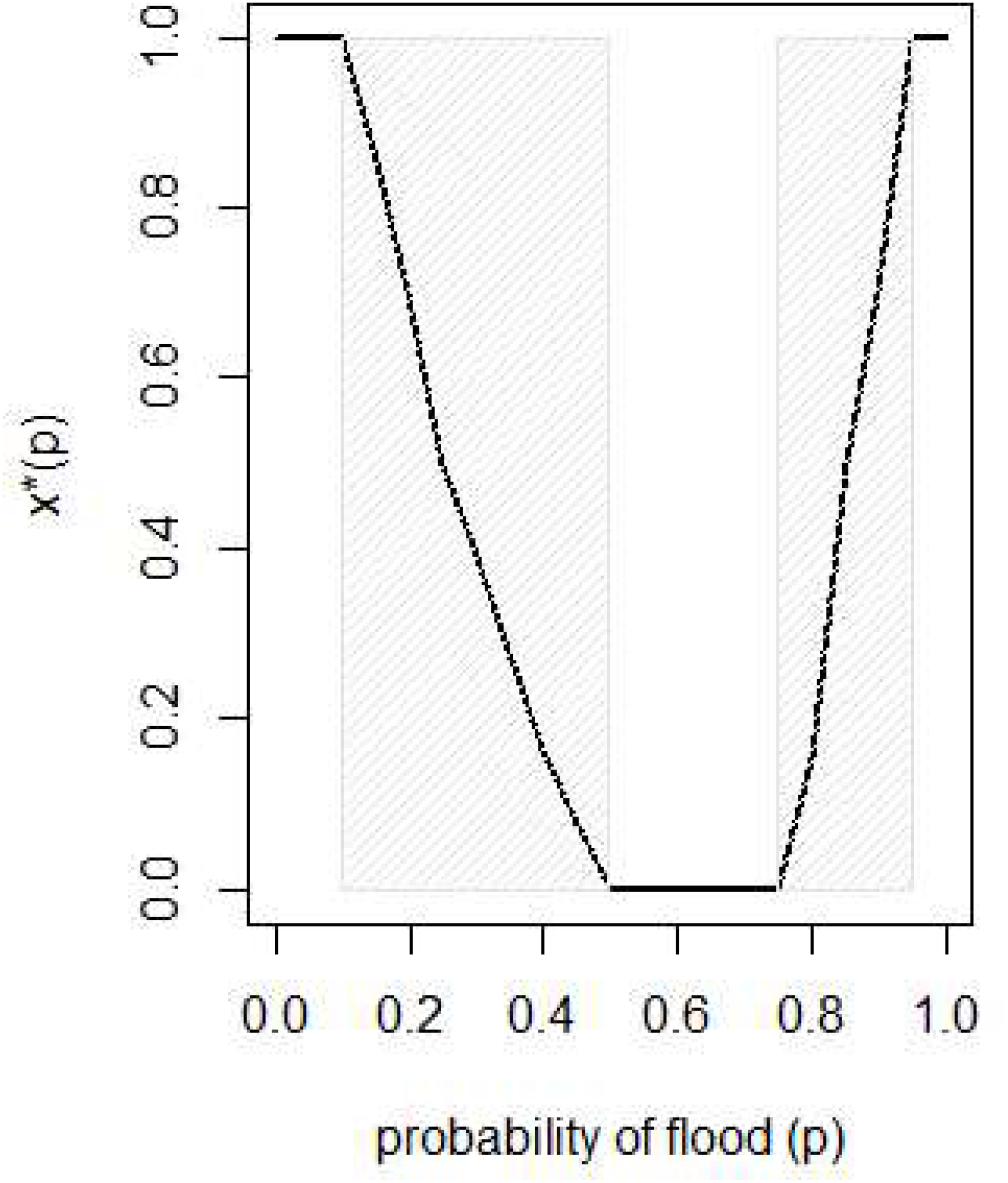

## An example with 4 environments and 2 phenotypes: drought and competition with other species in pioneer plants

In previous examples we represented our environments as a function of the frequency of one variable (p - the risk of floods) for simplicity. However, we showed in the section "Extension to finite populations" of Mafessoni, Lachmann and Gokhale that this is only a possible way to explore combinations of different environmental states (in Figure 5 shown as a potential trajectory in the simplex of potential combinations of environmental states).

Therefore, the probabilities of environmental states 1,2 and 3 might actually be set to take any probability values *p*_1_,*p*_2_ and *p*_3_ (with *p*_1_ + *p*_2_ + *p*_3_ = 1), to explore any ecological gradient of interest.

Here we show an hypotetical example in which multiple environmental factors affect the fitness of a species. Thus, the environmental states that our hypothetical species has to face are determined by the combinations of these factors. We take the example of rainforest pioneer plants. The growth of these plants is not only hindered by abiotic factors (for example water availability) but also by the presence of other plants. For this reason the seeds of these species, uniquely among other plants, break their dormancy in response to light. We now want to calculate the fitness benefit of informative cues (light and water) for the germination of an hypothetical species of rainforest pioneer plant. We assume that a seed of our pioneer plant might end up under the canopy (with probability p_canopy) or in a clear patch of forest (the latter being favorable), and that could have sufficient water (with probability p_wet) or not (p_arid=1-p_wet). Therefore we can have four possible environmental states (wet-canopy, wet-clear, arid-canopy, arid-clear) with probabilities p_wet x (1-p_canopy), p_wet x (1-p_canopy), (1-p_wet) x p_canopy and (1-p_wet) x p_canopy, respectively. This illustrates that interactions between multiple environmental factors can lead to asymmetric interaction matrices in which the number of potential environmental states exceeds that of phenotypes.

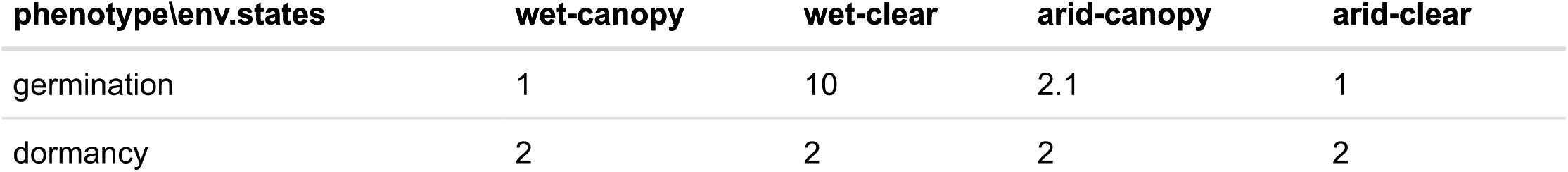

| phenotype\env.states | wet-canopy | wet-clear | arid-canopy | arid-clear |
| --- | --- | --- | --- | --- |
| germination | 1 | 10 | 2.1 | 1 |
| dormancy | 2 | 2 | 2 | 2 |

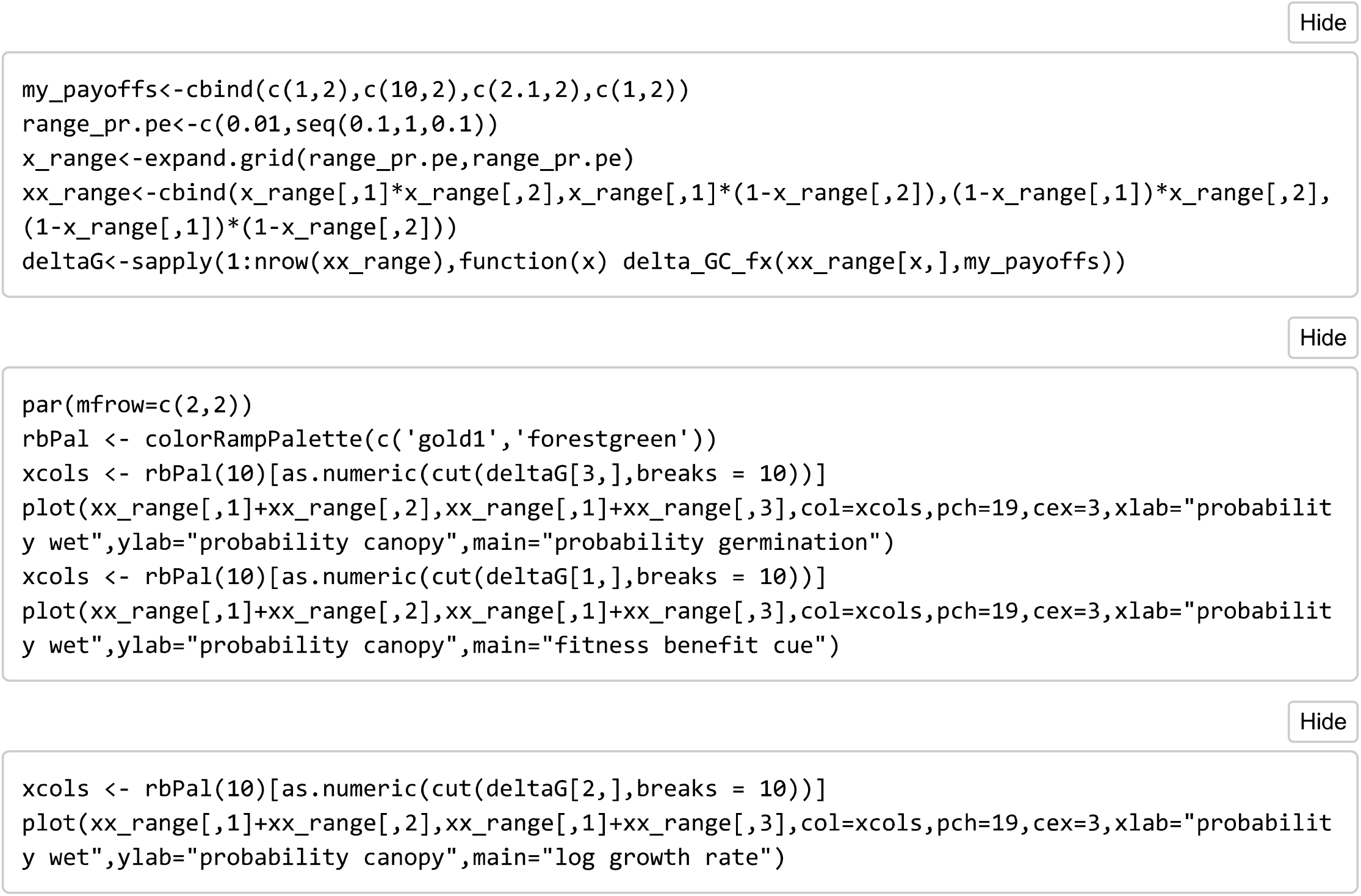

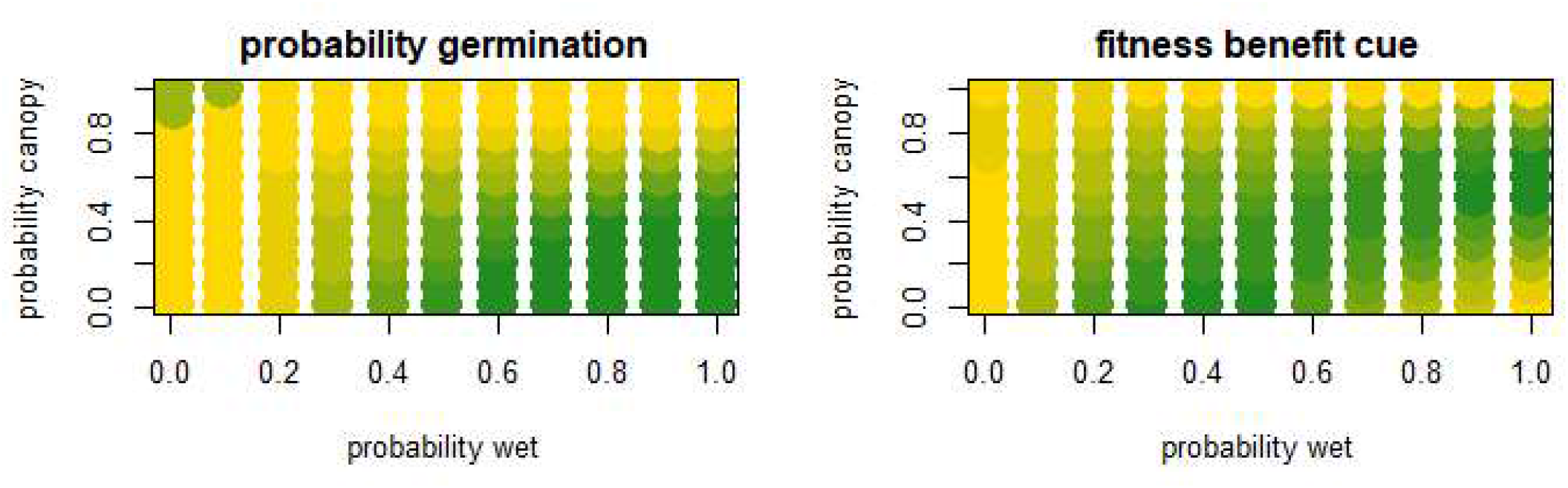

We can see that without cues plants would evolve to always germinate in clear areas, and hedge their bets in intermediate regions of aridity and canopy cover. For rainforest (high probability wet) the fitness benefit of a cue along the y axis reaches a peak for intermediate values. This indicates that plants adapted to light and wet conditions would have the highest fitness benefit in perceiving a cue about the canopy state (for example light) for intermediate covers of canopy. A possible biological prediction is that plants whose seeds respond to light to interrupt dormancy are unlikely to evolve in forest that show very few clear patches or that are very sparse. Conversely, we see that along the x-axis (probability of wet), for small values of canopy covers (plants that live in open areas), the fitness benefit of a cue is maximum for intermediate values of wetness-aridity.

## The fitness value of perceiving lunar emergence and tides in hypotetical intertidal organisms

The reproductive behaviour of many marine organisms living in intertidal environments is strongly influenced by tidal patterns. A correct timing for the emergence of the adult individuals of these species is critical in natural populations, since high tides can wash away the individuals, preventing their reproduction. For example, a favorable time for the reproduction of the marine midge *Clunio marinus* (Chironomidae, Diptera) occurs only for about 4 hours in a lunar cycle. *Clunio* individual rely on a complex a circalunar clock to obtain cues on when to emerge. Here we sketch an hypothetical estimate of the theoretical fitness advantage of the complex adaptative mechanism of similar organisms. We stress the fact that \***these calculations are purely hypothetical**, *as currently not enough data is available to accurately estimate the fitness consequences of different scenarios for this species, and only approximate calculations can be made. Therefore We denote our hypotethical species simply as* moonlight*.

To illustrate this case, we model it in two different ways. In both cases we rely on the intuition that we can represent our system in terms of two different environmental states (low tide or high tide). The frequency of low­ tides, which are apt for reproduction, is: frequency of conditions apt for reproduction ~ 4/(28*24) ~ 0.6%

First, we could consider a 2×2 environment-behavior matrix with two phenotypes, dormancy and emergence, similarly to the cases of annual plants described above. Every hour individuals can potentially emerge or not. In absence of any cues, *moonlight* individuals could hedge their bets whether to emerge and face the risk of encountering a high-tide, or wait one more hour. Since an individual cannot reproduce if the tide is high, we assume payoff O in that case. When an individual emerges at low-tide manage, its payoffs is *n_1_*, which we can imagine as the number of larvae passed to the next generation. Dormant individuals might emerge later. Since we do not know how long *moonlight* could survive without emerging, we specify a payoff for dormancy of 1 - 1 / (24 * 28), which can be seen as the probability that an individual survive to the next hour assuming an average lifespan of one lunar cycle and a constant death rate. In absence of cues:

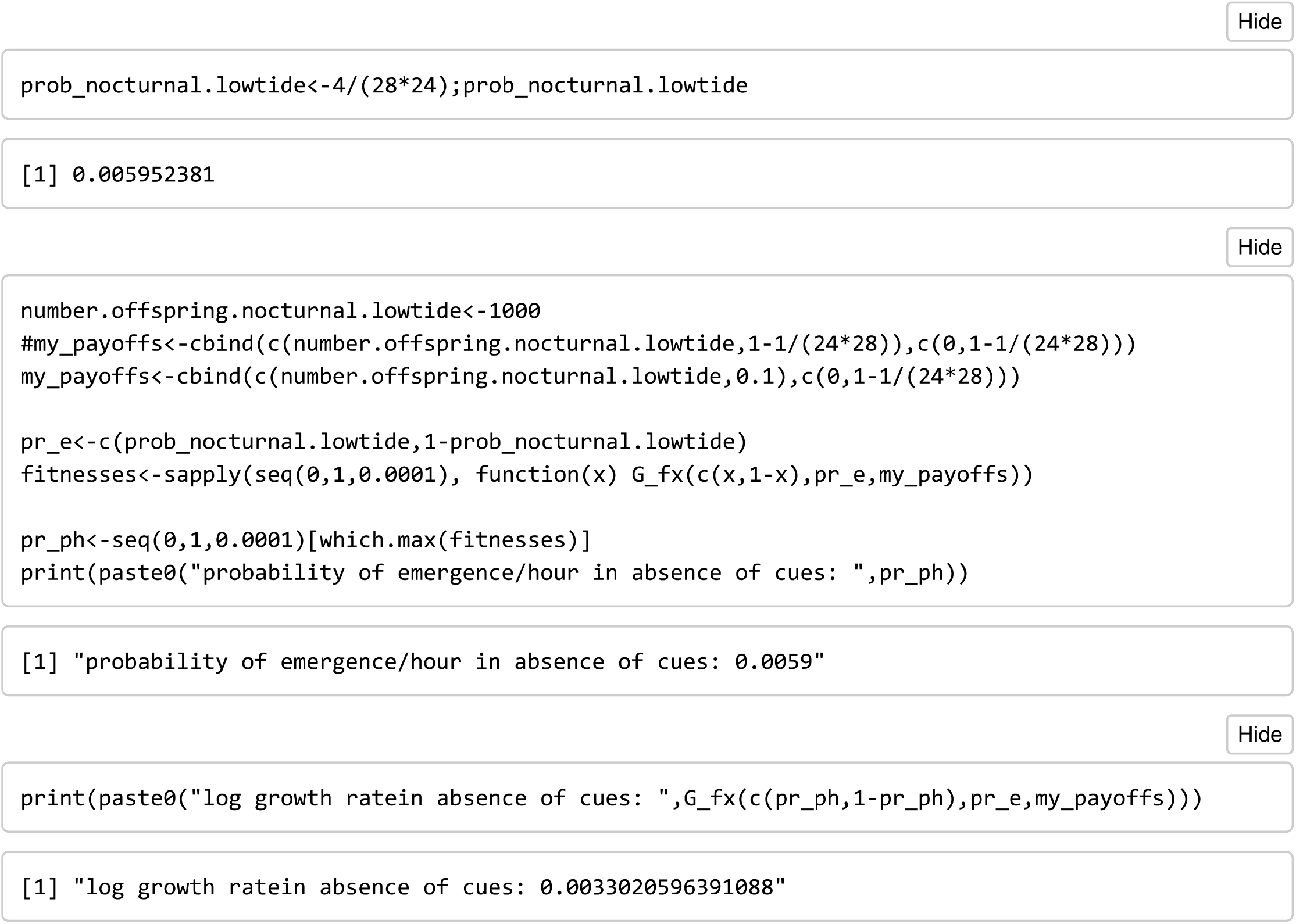

When informative cues are used instead the log growth rate is:

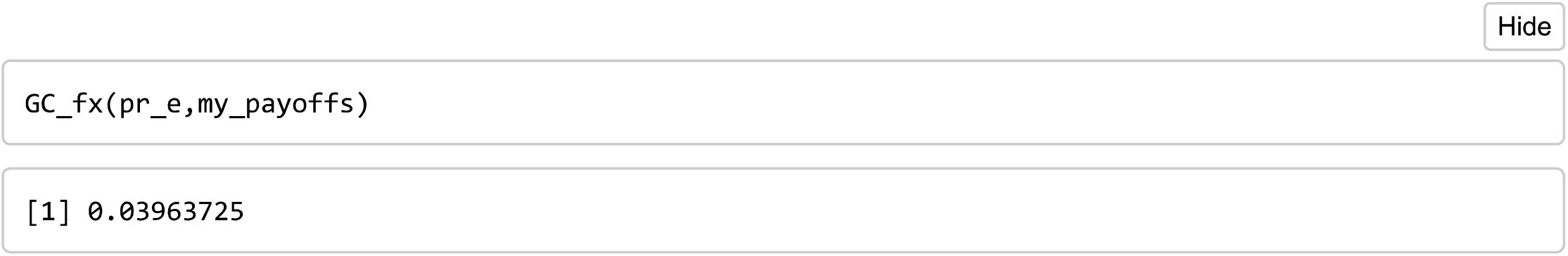

Hence, the fitness value of the complex adaptative strategy of Clunio is (in terms of bits):

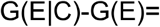

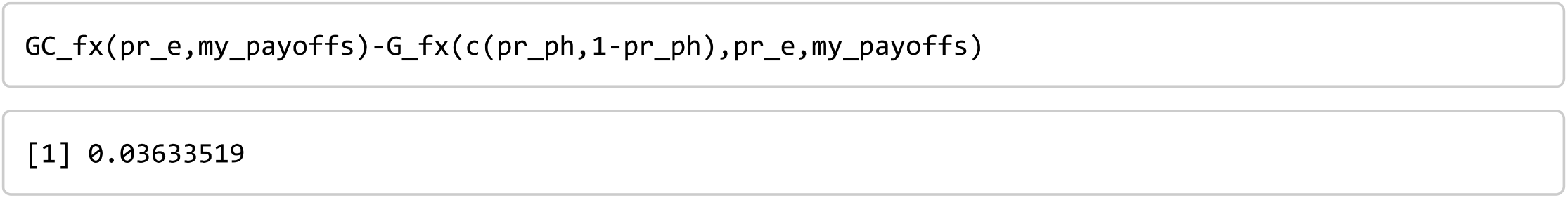

Notice that here we made some assumptions about the life cycle of Clunie. For example we assumed that each successful! reproduction event might lead to 1000 larvae. How does the fitness benefit depend on the number of larvae?

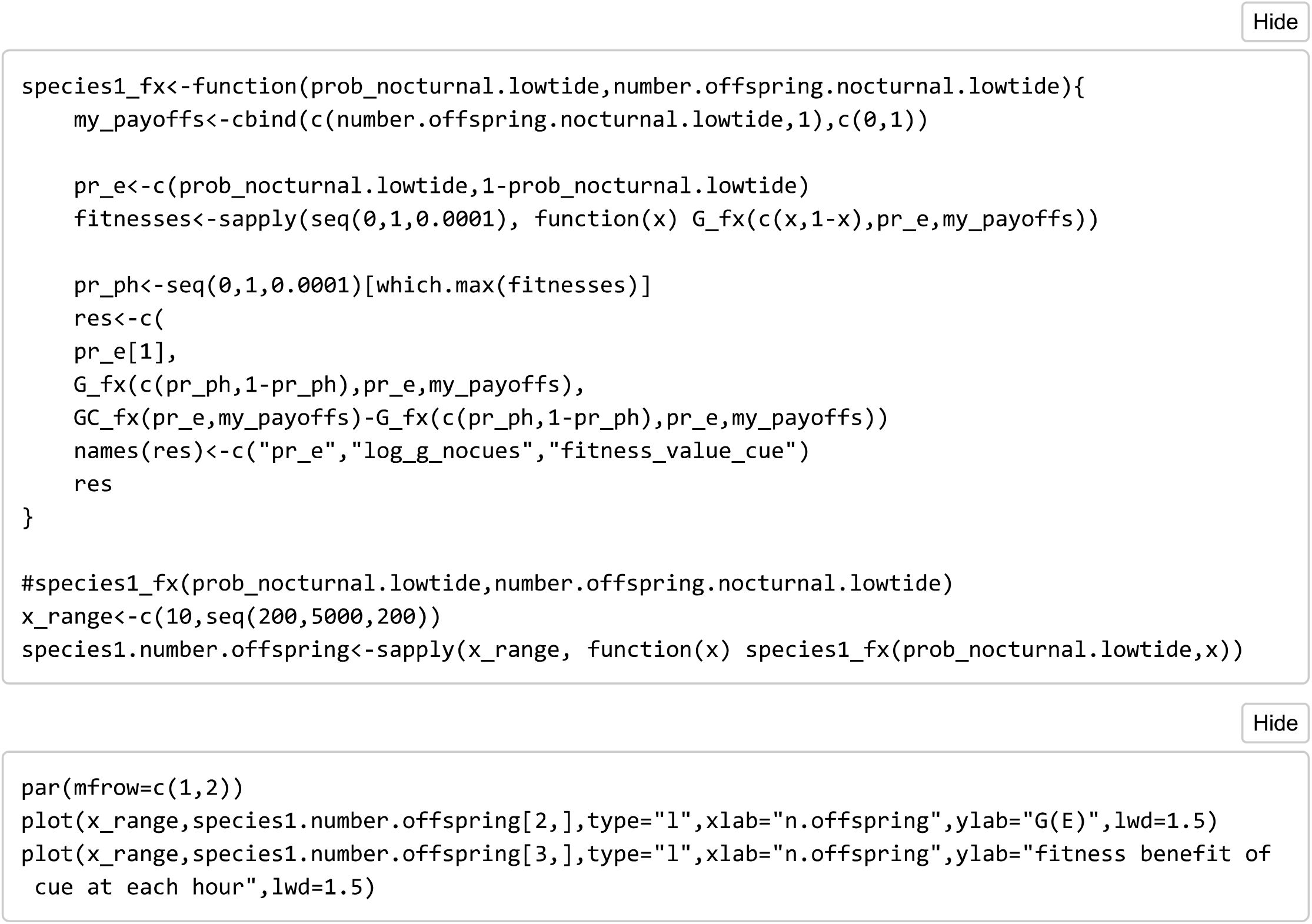

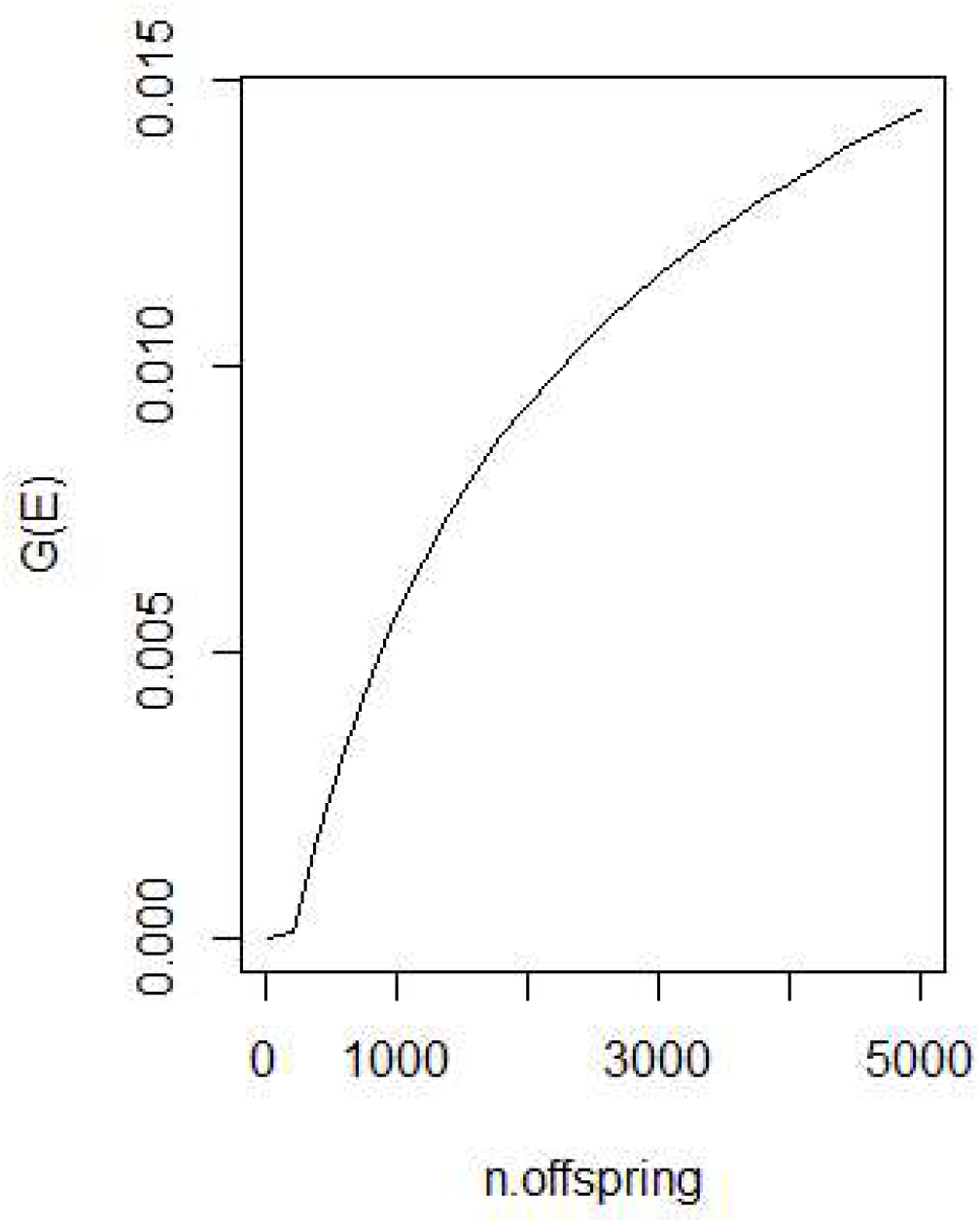

Note that here the fitness value of a cue is considered for an individual time unit (here hours) of the life cycle of a species of *moonlight.* To obtain the value for the entire cycle, we just have to multiplicate by its lenght. So, for life­ cycle of approximately 28 days, **a clue for a *moonlight* with many larvae would have approximately 5 bits of value**. Interestingly, different populations of Clunio in different geographical regions have been found with different circalunar and circadian patterns. Specifically, in some populations Clunio emerges exclusively with full moon, while in others exclusively with the new moon low-tide. In these cases adults can achieve sexual reproduction only in one tide-light condition per lunar cycle, most often with the new-moon.

However, in some populations individuals bet-hedge between the two lunar phases in equal proportions. In these cases, two potential reproductive time windows exist.

How does the fitness value of the lunar signal differ in the two cases? In the plots we can see G(E) and the fitness value of an informative cue about the circalunar phase, for a population all synchronously emerging at the new­ moon low-tide (1 m), or populations in which individuals can emerge either in new or full moon low-tides (2m). At low proportions of favorable tide-light conditions for reproduction the increase in fitness value increases roughly linear.

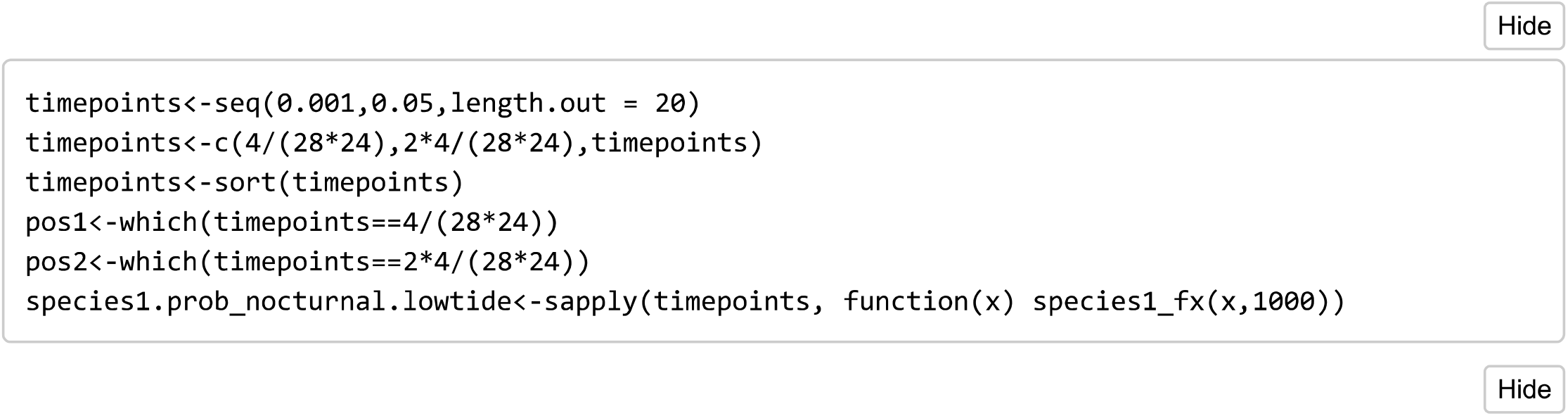

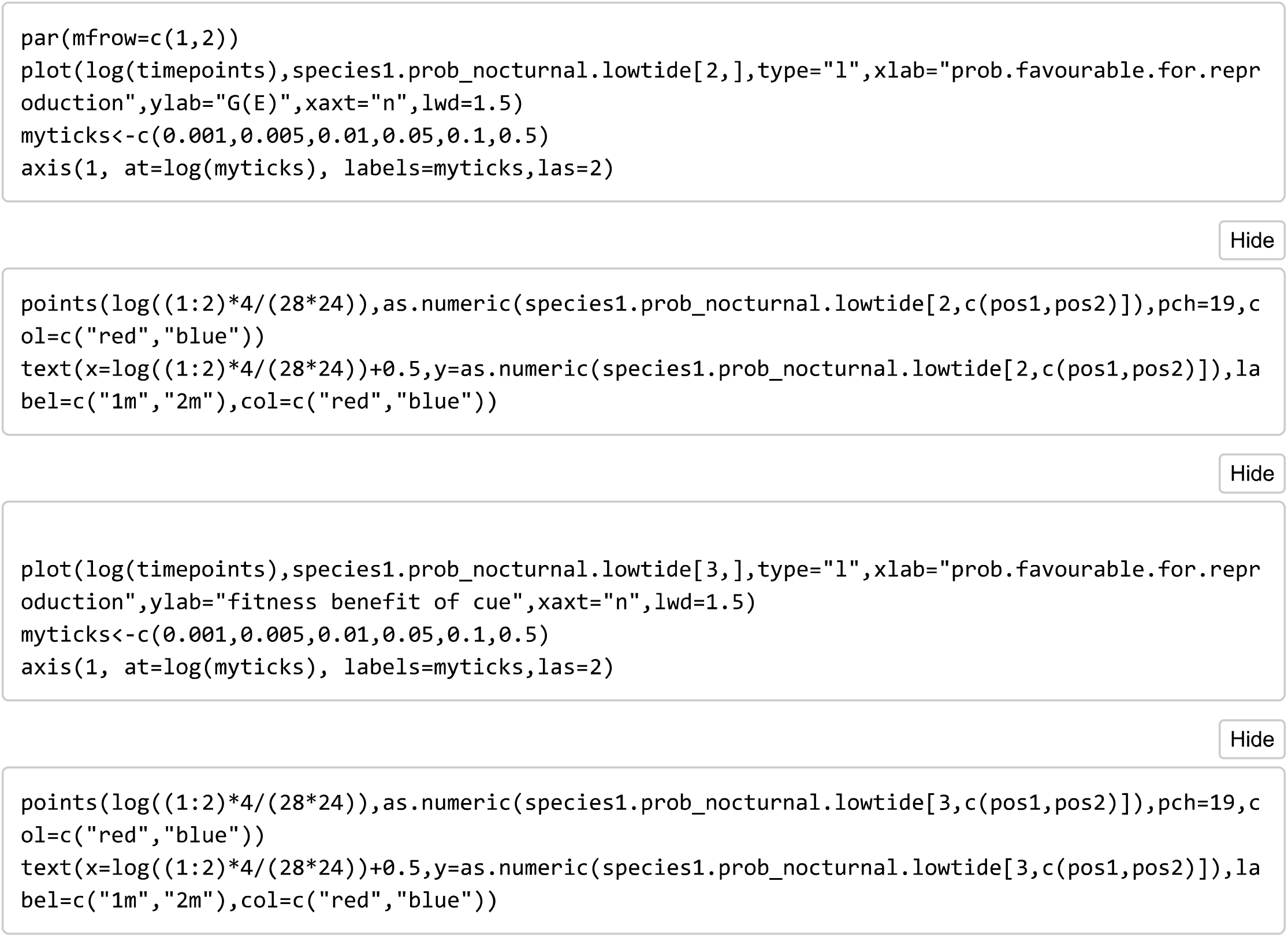

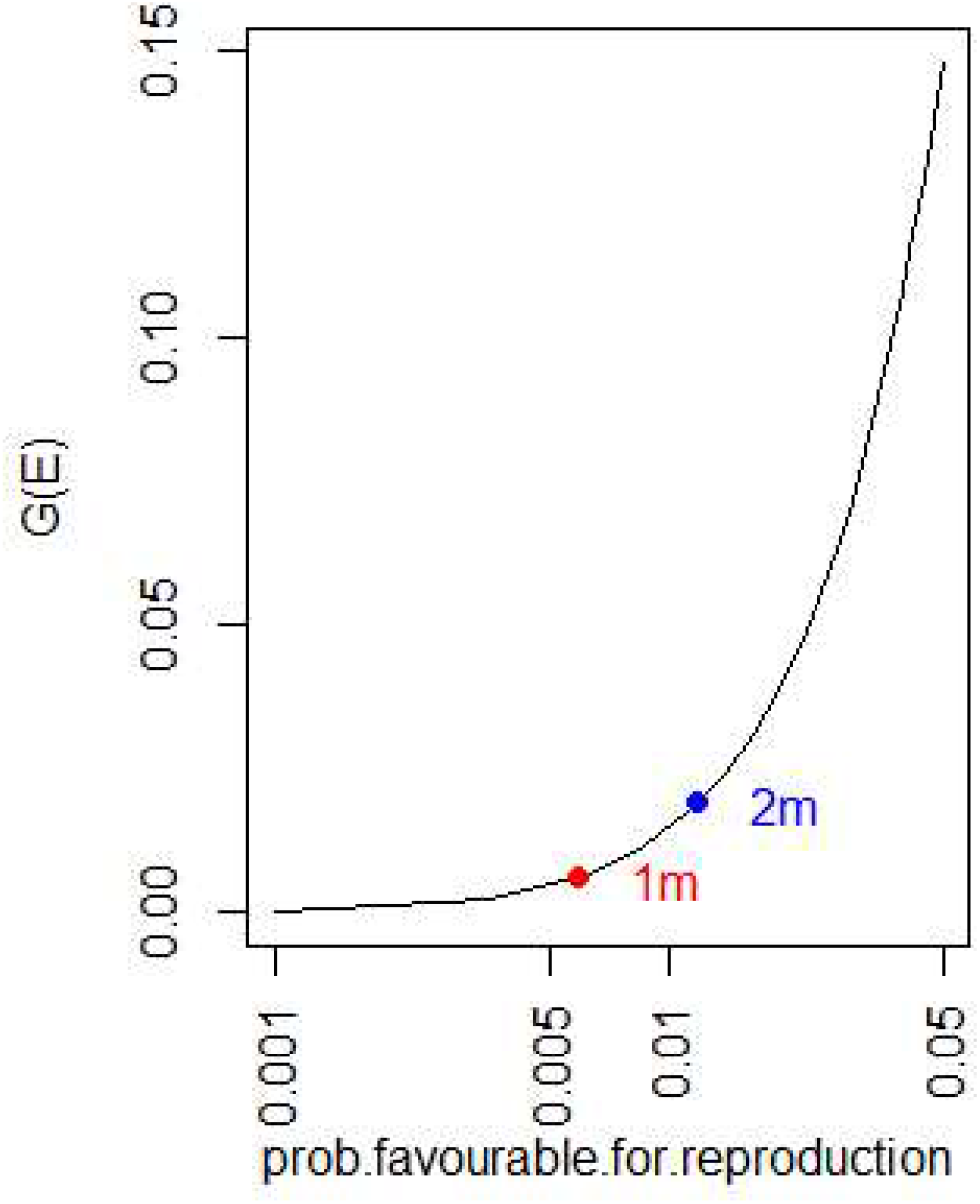

There are some problems with this model: first, we do not know the payoffs second, it might be inappropriate to assume a constant death rate; third, we might want to model explicitly different strategies (which have for example been documented in *Clunio)* and cues; last, we know that in a month there is always a fixed number of low-tides.

To circumvent these issues, we can think of a simple second model, in which the possible phenotypes are the different circalunar and circadian conditions for adult emergence. In a lunar month there are 28 days (28 lunar phases) and for each day several time-light conditions. Since adults live 2-4 hours we consider 6 4-hour long time­ light conditions each day, for a total of 168 moon-light conditions. Individuals can hedge their bets across them.

Since reproduction cannot occur if adults do not emerge during low tides we have a proportional bet-hedging case, we can collpase the 168×168 matrix to a 2×2 matrix:

| emerge time | time1 | time2 | ... | time168 |
| --- | --- | --- | --- | --- |
| emerge time 1 | 1 | 0 | 0 | 0 |
| emerge time 2 | 0 | 1 | 0 | 0 |
| .. | .. | .. | .. | .. |
| emerge time 168 | 0 | 0 | 0 | 1 |

where we define time1 the actual time of low tides.

In absence of cues the probability of emergence is (since we have proportional bet-hedging) 1/168, and log growth rate is *log*(l/168).

When informative cues are used instead the log growth rate is simply O (if the payoff is 1 to reflect a population at the steady state).

Hence, the fitness value of the complex adaptative strategy of Clunio is (in terms of bits): G(E|C)−G(E)=

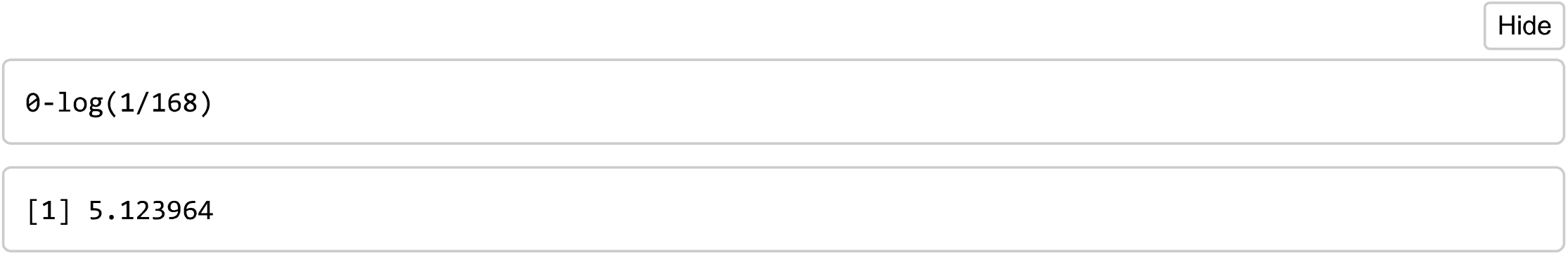

which is similar to what obtained above for large offspring sizes. Note that we can also look separately at the value of the information coming from the lunar cycle and circadian one. An organism relying only on the moon cycle has still no clue at what time of the day to emerge - since this is informed by the light cycle. Hence, an organism would hedge its bets across the 6 time of the day conditions. Its log growth in absence of light information will be:

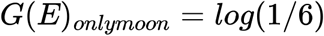

while one not being able to rely on the lunar cycle but only on time-light:

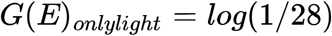

Hence, the fitness value of lunar and circadian clues are:

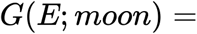

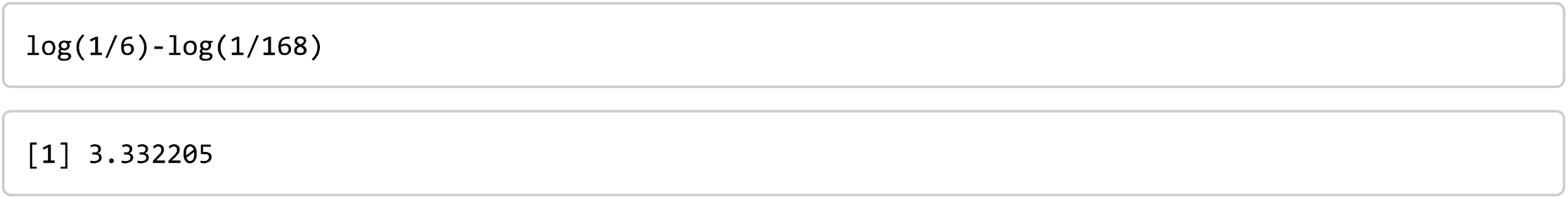

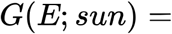

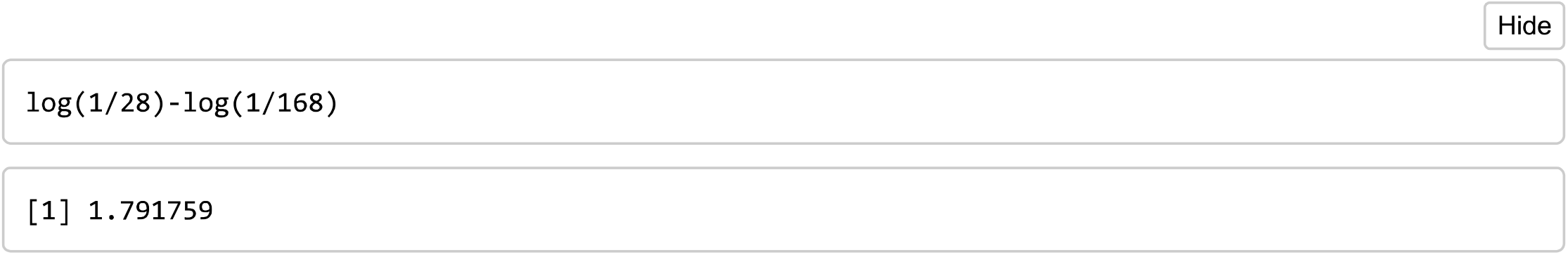

indicating that the lunar cycle is about twice as important as time cues in terms of fitness for *moonlight*.

